# Linear modeling reveals a predominance of *cis*- over *trans*- regulatory effects in wild and domesticated barley

**DOI:** 10.1101/661926

**Authors:** Matthew Haas, Axel Himmelbach, Martin Mascher

## Abstract

Barley, like other crops, has experienced a series of genetic changes that have impacted its architecture and growth habit to suit the needs of humans, termed the domestication syndrome. Domestication also resulted in a concomitant bottleneck that reduced sequence diversity in genes and regulatory regions. Little is known about regulatory changes resulting from domestication in barley. We used RNA-seq to examine allele-specific expression (ASE) in hybrids between wild and domesticated barley. Our results show that most genes have conserved regulation. In contrast to studies of allele specific expression in interspecific hybrids, we find almost a complete absence of *trans* effects. We also find that *cis* regulation is largely stable in response to short-term cold stress. Our study has practical implications for crop improvement using wild relatives. Genes regulated in *cis* are more likely to be expressed in a new genetic background at the same level as in their native background.

## Introduction

Barley (*Hordeum vulgare* ssp. *vulgare* L.) is an important crop for feed, malting and to a lesser extent, human consumption (Ullrich 2010). Among the first crops to be domesticated in the Fertile Crescent about 10,000 years ago (Zohary et al. 2012), barley remains fully interfertile with its wild progenitor *H. vulgare* ssp. *spontaneum* K. Koch (*H. spontaneum* for short). Therefore, *H. spontaneum* is considered to be a useful source of beneficial alleles for barley improvement. Preferential selection of genotypes with traits beneficial to humans and the intentional breeding have narrowed the genetic diversity and altered gene expression patterns. These molecular changes have caused differences in plant architecture and growth habit between wild and domesticated relatives, collectively called the domestication syndrome (Hammer 1984; Doebley et al. 2006).

In barley, key domestication and crop evolution genes include *Non-brittle rachis 1* (*btr1*) and *Non-brittle rachis 2* (*btr2*) controlling dehiscence of spikelets from the rachis; *six-rowed spike 1* (*vrs1*), which is responsible for lateral floret fertility and may be modified by *INTERMEDIUM-C* (*INT-C*); *VERNALIZATION1* (*Vrn1*) which controls the vernalization requirement; covered/naked caryopsis *(nud*) affecting the adherence of the hull to the caryopsis; and *Photoperiod-H1* (*Ppd-H1*) affecting photoperiod sensitivity (Trevaskis et al. 2003; Turner et al. 2005; Komatsuda et al. 2007; Taketa et al. 2008; Ramsay et al. 2011; Pourkheirandish et al. 2015). These genes were cloned using traditional mapping approaches as their effects are easy to observe given the major phenotypic effect of each gene; however, these tasks were also facilitated by the relative ease for which DNA sequence variation is detected between unrelated genotypes. The task of detecting regulatory variation is more challenging since DNA sequence data alone cannot be used to predict expression. Regulatory variation may arise due to differences in *cis* or *trans* factors. *Cis* factors are physically linked to the genes they control such as promoters or enhancers while *trans* factors act distally, such as transcription factors. Many studies have been conducted to study the effect of domestication on gene regulation (Rapp et al. 2010; Swanson-Wagner et al. 2012; Koenig et al. 2013), although these studies were not designed to disentangle *cis* and *trans* effects.

In order to achieve separation of *cis-* and *trans-* acting factors, Cowles et al. (2002) proposed the comparison of allele-specific expression (ASE) in F_1_ hybrids to that of the parents. Subsequently, Wittkopp et al. (2004) demonstrated how to find the relative contribution of *cis* and *trans* factors. We show this in Supplementary Figure S1 and provide further explanation in the Methods section. Zhang and Borevitz (2009) conducted a similar study using a custom gene expression array with allele-specific probes; however arrays are known to suffer from ascertainment bias (Nielsen 2000). In addition, it can be challenging to design suitable probes that can distinguish between two alleles as demonstrated in yeast by Tirosh et al. (2009). The advent of low-cost RNA-seq enabled the strategy of genome-wide total and ASE to be implemented in a single experiment in *Drosophila* (McManus et al. 2010). Lemmon et al. (2014) extended this approach to examine regulatory changes between maize and its wild progenitor, teosinte. Cubillos et al. (2014) used the approach to examine the steady-state stress drought response in *Arabidopsis*. To the best of our knowledge, only one previous study has been published examining ASE in barley (von Korff et al. 2009). In that study, the authors used custom gene expression arrays to measure ASE ratios for 30 stress-response genes in five F_1_ hybrids at different developmental stages. In the present study, we used RNA-seq to estimate the impact of domestication on gene regulation in barley and whether the response to an environmental stress (cold) is affected by domestication.

## Results

### Experimental design

The experimental design is summarized in Figure 1. Plants were grown in duplicated trays for one week in a growth chamber (22°C day/18°C night) with a 12 h photoperiod. On the day of the cold treatment, one of the trays was moved to a vernalization chamber (4°C) for three hours (11:00-14:00). The cold treatment and tissue harvesting were done at the same time of each day to avoid confounding factors due to circadian rhythm. The experiment was conducted four times. A fifth replicate was added in order to get additional replicates for samples which failed during the previous four attempts. For randomization, the layout of plants in the trays was changed for each replicate, but both trays within a replicate had identical layouts.

**Figure 1.**
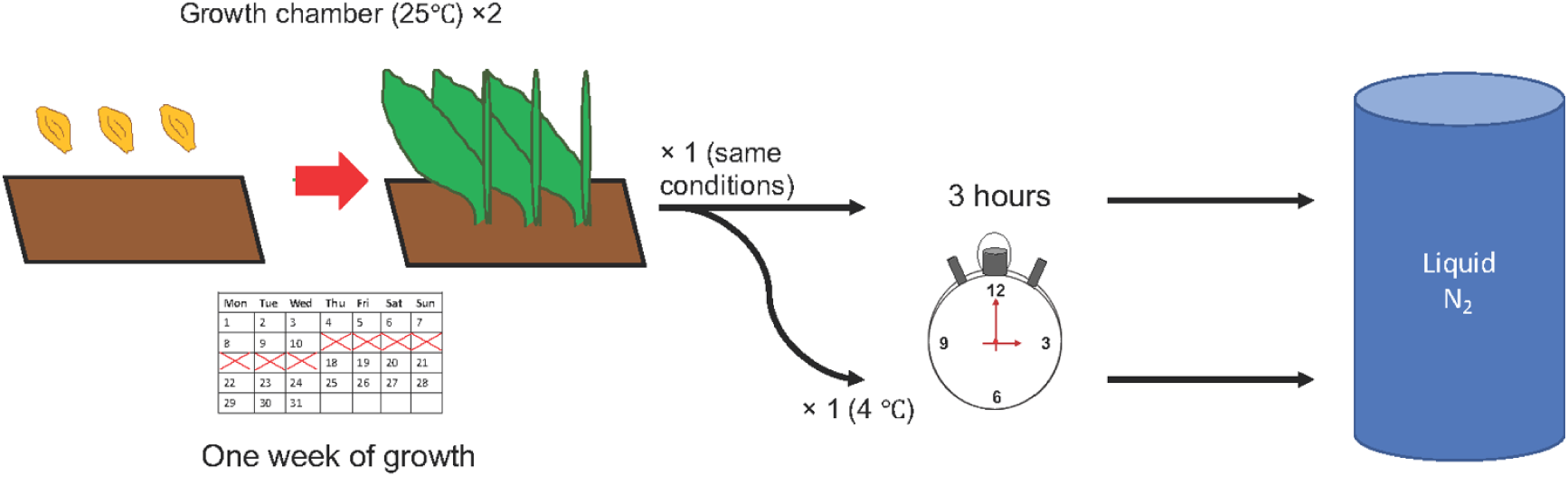
The experimental design. Barley seedlings were grown for one week until the first leaf was fully expanded. in duplicated trays. After one week of growth, one tray was moved to a cold room (4°C) for three hours (from 11:00-14:00) while the other tray remained in the growth chamber (22 °C). After three hours, samples were collected in liquid nitrogen and stored at −80°C until we prepared them for RNA extraction.

### Plant material

Three cultivars, two landraces and four wild accessions were used in this study for a total of nine parental lines (Table 1). This includes the common maternal reference, Morex (CIho 15773), a six-rowed spring malting cultivar from Minnesota, USA (Rasmusson and Wilcoxson 1979). All other accessions were crossed to Morex, bringing the total number of genotypes to seventeen. Morex was selected because the recently released barley reference genome was generated from bacterial artificial chromosome (BAC) sequences originating from this cultivar (Mascher et al. 2017). The other accessions were selected from an exome capture panel of 267 wild and domesticated barleys in order to maximize geographic and genetic diversity (Russell et al. 2016). Principal component analysis (PCA) of 1.7 million bi-allelic SNPs from these data separate wild samples based geography and domesticated samples based on breeding history as well as row type (Figure S2). The target space is 60 Mb or about 75% of the barley gene space (Mascher et al. 2013). Each parental genotype was previously subjected to at least two rounds of single-seed descent to decrease residual heterozygosity.

**Table 1.**
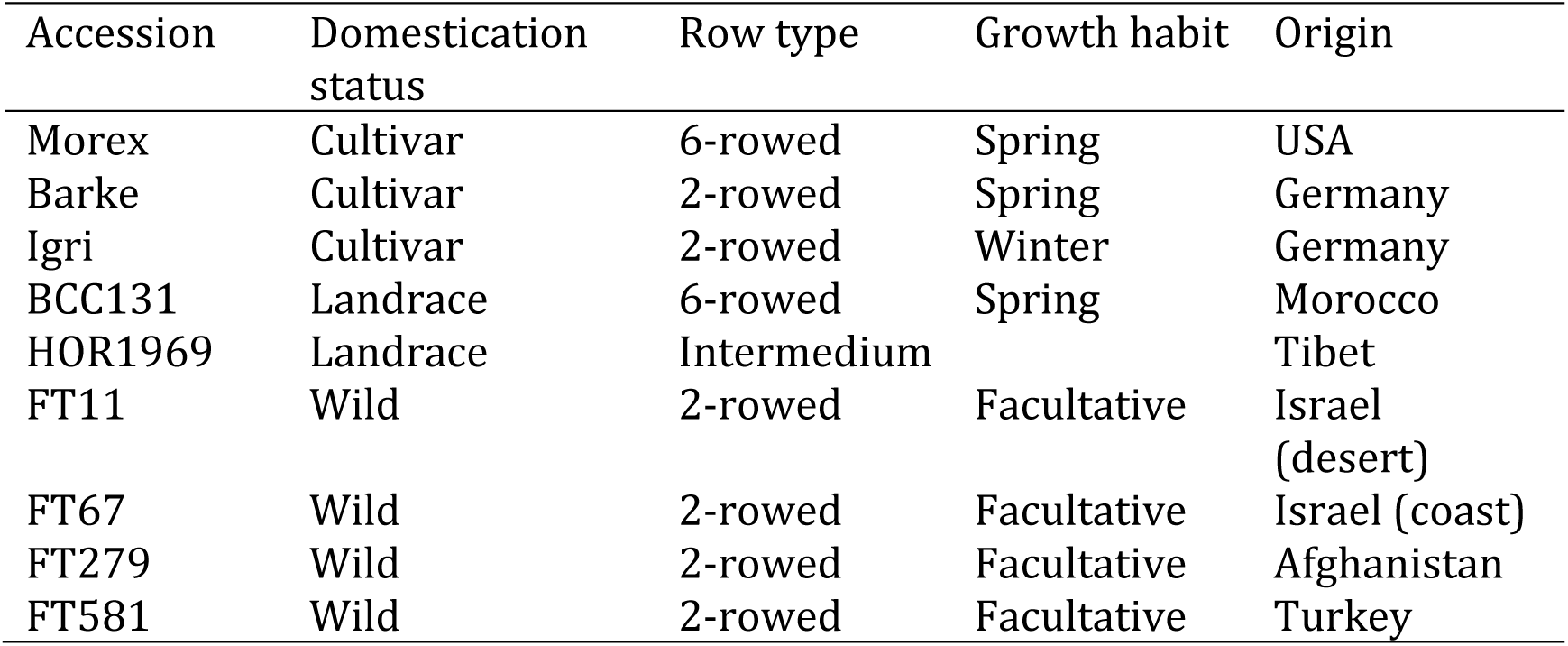
Accessions used in this research.

### Data quality

Most samples mapped to the barley reference sequence at a high rate (>= 80%), but eight samples (all from genotype BCC131) had a mapping rate of less than 80% (Figure S3-S4).. Six of these had a mapping rate between 50-79% and one sample had a mapping rate of 30%. To determine the cause of the low mapping rate of the eight samples, a Basic Local Alignment Search Tool (BLAST) run was conducted. For those samples with a mapping rate between 50-79%, the source of contamination is the barley stripe mosaic virus (BSMV, Figure S4) while the sample with the lowest mapping rate (30%) is contaminated with human DNA (Figure S5). BCC131 samples were included in our analyses anyway because the effect of sequence contamination, reduced coverage, merely reduces statistical power for variant calling. While this reduction decreases power for ASE and differential expression analysis, the data for genes that remain informative are still useful.

### Principal component analysis

After checking our gene expression data quality, we examined the data to see if it matches our expectations to ensure that it is reliable. Principal component analysis was conducted using kallisto-derived expression data. The first principal component explains 25% of the variance and separates the parental genotypes from Morex, the common maternal parent for all hybrids. The hybrids cluster between Morex and the parents, as expected for hybrids (Figure 2A). The second principal component explains 9% of the variance. Three parental samples (BCC131, Barke and Igri) form a cluster separate from the other accessions. The best explanation for this is geography. The cultivars Barke and Igri are from Germany and BCC131 is a Moroccan landrace, while wild barleys FT11 and FT67 originate from different environments in Israel, FT279 is from Afghanistan, FT581 is from Turkey, and the landrace HOR1969 is from Tibet (Figure 2B). The third principal component explains 8% of the variance. Samples along this component cluster by accession; however, only HOR1969 loosely clusters separately from the others (Figure S6). The fourth principal component explains 7% of the variance and separates samples according to treatment (Figure 2C). The PCA results show that samples cluster according to the principal factors in our experiment (i.e., generation, genotype and treatment). Therefore, the data may be used to confidently determine allele-specific expression.

**Figure 2.**
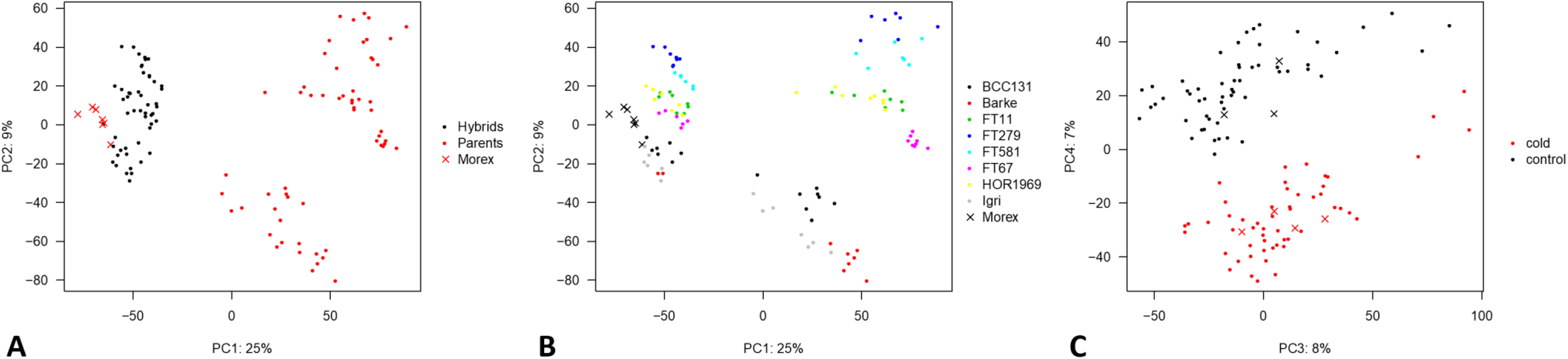
Principal component analysis: (A) PC1 separates samples based on generation. Parental samples cluster on the right, while hybrids cluster on the left, closer to the common reference parent, Morex. (B) PC2 separates samples based on accession. Accessions broadly cluster by geography. (C) PC4 separates samples according to treatment.

### Exome capture and SNP calling

The PCA described above was conducted using kallisto, but these results are for overall expression and are not allele-specific. HISAT2 was used for allele-specific mapping of reads. In order to determine which allele a transcript originated from, exome capture data were collected for one individual of each hybrid. Exome capture and (in some cases) whole genome shotgun data already exist for the parents. By comparing SNPs between parental accessions and confirming these SNPs in hybrids between these accessions and transcript (RNA) data, we were able to unambiguously assign transcripts to one allele or the other. This section describes in detail how we found these SNPs. Variant call format (VCF) files resulting from the exome capture analysis pipeline were more rigorously filtered in R. Coverage filters were applied to produce a set of high-confidence SNPs for each cross combination. These filters required a minimum quality score of 5 for both homozygous and heterozygous genotype calls, a minimum read depth of at least 10 for both homozygous and heterozygous genotype calls. The minimum fraction of heterozygous call was set to 1 since we were working with hybrids. The maximum fraction of missing genotype calls was set to 0.9 and the minimum minor allele frequency was 0.05. Genomic Data Structure (GDS) files were produced using the R package SeqArray (Zheng et al. 2017) which contain clear differences between reference and alternate alleles. Exome capture mapping statistics are presented in Figure S7. The number of informative SNPs and genes are presented in Table 2. SNPs are informative if they reside in genic regions since SNPs are only useful for ASE when they are transcribed. SNPs in regulatory regions are important for ASE, but they cannot be detected from RNA-seq data. The number of informative genes for BCC131 (2,589) is lower than expected based on the other landrace, HOR1969 (6,850 genes) as a result of lower coverage due to contamination as discussed above (Figures S3–S5). Otherwise, the general pattern of wild accessions being more diverged from Morex (8,282 – 9,318 informative genes) compared to cultivars (4,296 and 4,634 genes for Barke and Igri, respectively) is, unsurprisingly, observed.

**Table 2.**
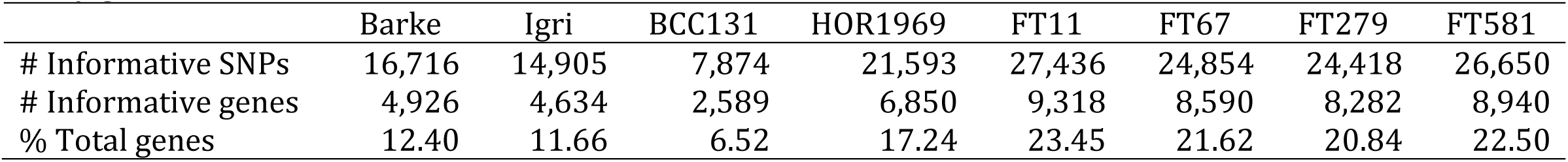
The number of informative SNPs, the number of informative genes and the percent (%) of total high-confidence genes in the barley genome between each accession relative to the cultivar Morex.

|  | Barke | Igri | BCC131 | HOR1969 | FT11 | FT67 | FT279 | FT581 |
| --- | --- | --- | --- | --- | --- | --- | --- | --- |
| # Informative SNPs | 16,716 | 14,905 | 7,874 | 21,593 | 27,436 | 24,854 | 24,418 | 26,650 |
| # Informative genes | 4,926 | 4,634 | 2,589 | 6,850 | 9,318 | 8,590 | 8,282 | 8,940 |
| % Total genes | 12.40 | 11.66 | 6.52 | 17.24 | 23.45 | 21.62 | 20.84 | 22.50 |

### Assignment of genes to regulatory categories

For each of the informative genes, we mapped transcripts to determine whether or not there was allele-specific expression. Initially, we followed the methods used by McManus et al. (2010); however, as we inspected expression plots further, we realized that genes assigned to the *trans* only category differed greatly in their expression levels between replicates (Figure 3B). Use of the linear model resulted in a drastic reduction in the number of genes with *trans* effects including *trans only*, *cis + trans* and *cis × trans* (Figure 4A, Table 3) compared with the binomial method used by (McManus et al. 2010)(Figure 4B, Table 4). This is in line with what other authors have found in other organisms (Goncalves et al. 2012; Osada et al. 2017). Another notable trend is that the number of genes assigned to the conserved class of regulatory variation is higher when using a linear model. Approximately 80% of the total number of genes were assigned to this class using a linear model versus ∼20% using the binomial/Fisher’s exact test (Tables 3-4). In addition, regulation of gene expression appears to be stable in response to environmental stress, consistent with the findings of Cubillos et al. (2014). Regulatory category plots for all crosses are given in Figure S8.

**Figure 3.**
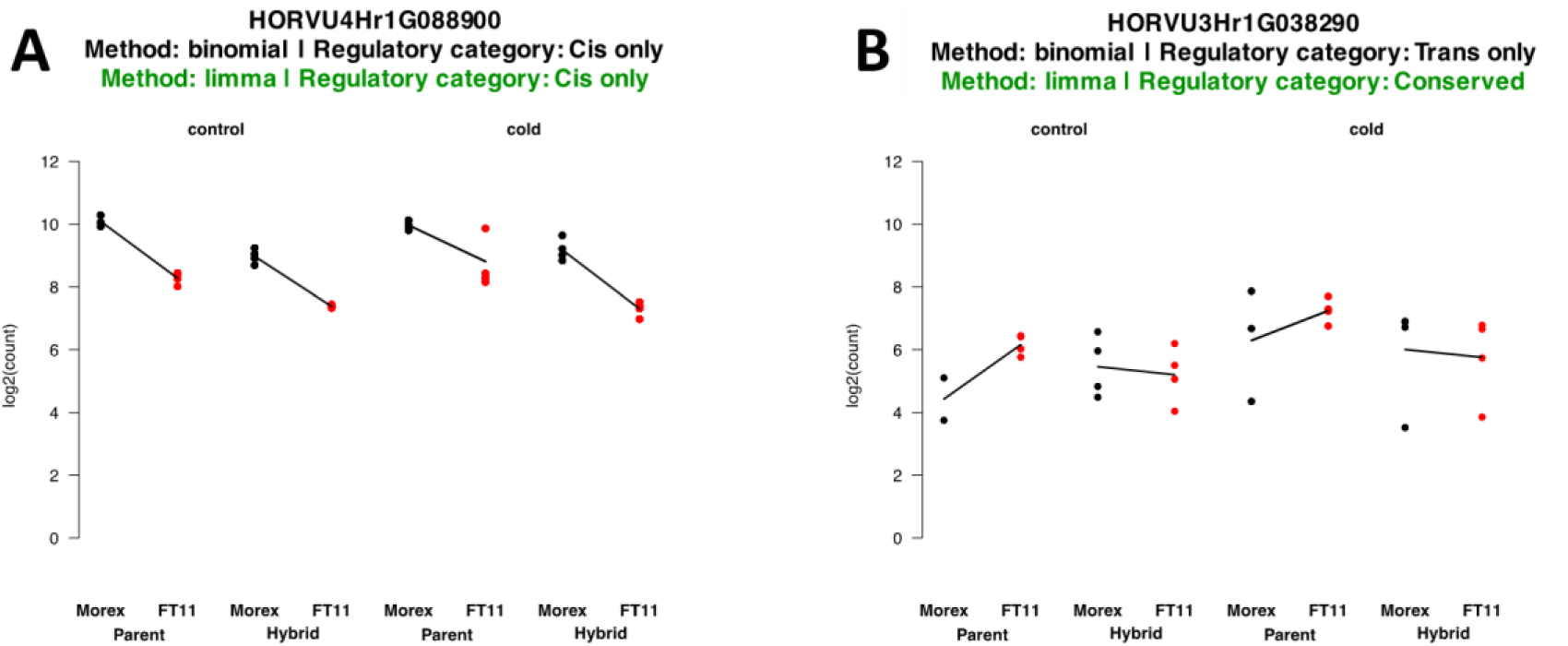
Example profiles for two genes illustrate the effect of the statistical differences between the binomial testing and linear model methods. Both methods agree in *A* because of the similar expression values between replicates; however, in *B* the large differences in expression between replicates mean that confidence in the true expression value is low.

**Figure 4.**
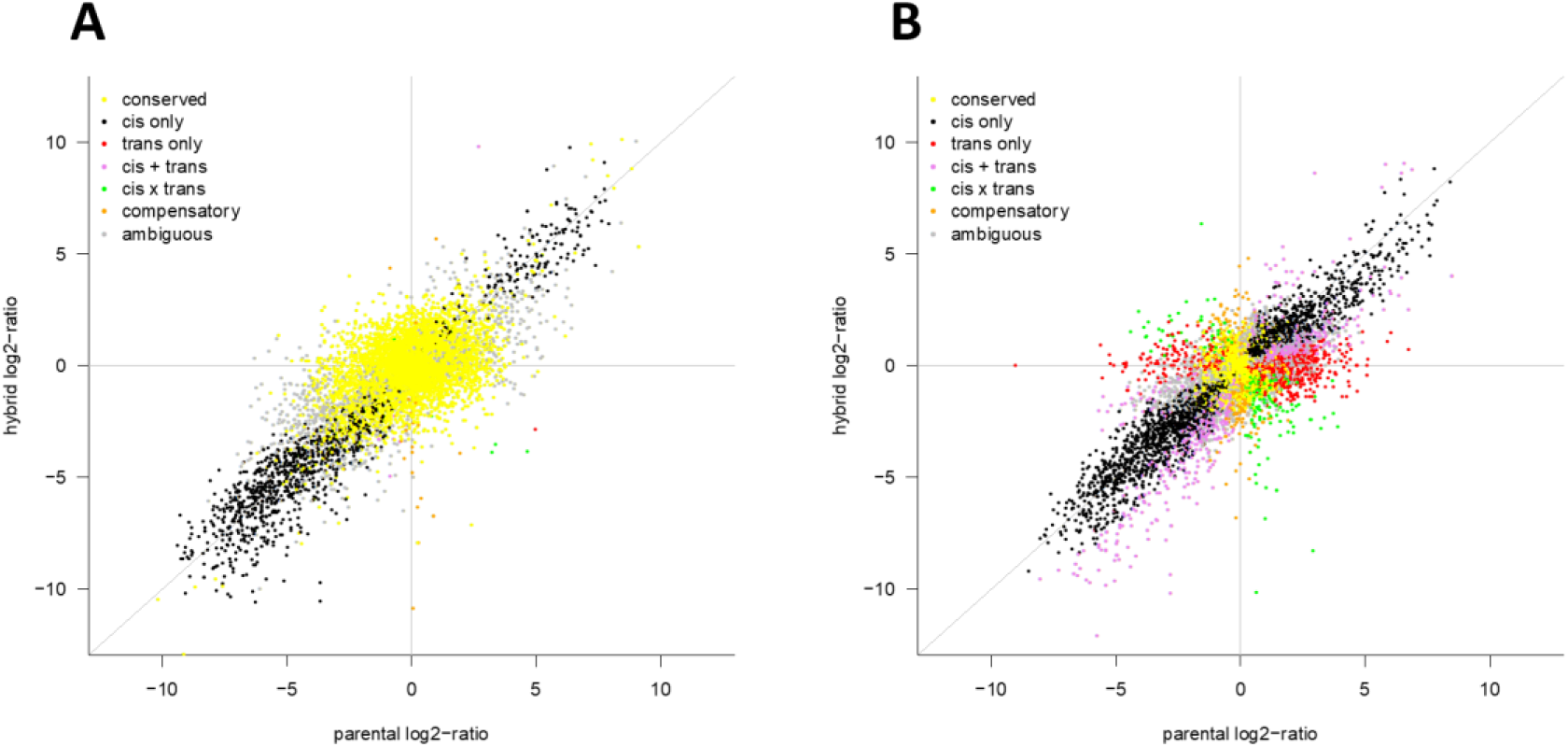
A global view of regulatory category assignment for one hybrid and its parents, in this case Morex and FT11. The x-axis shows the log_2_ ratio of expression difference between the parents, while the y-axis shows the log_2_ ratio of expression difference between the parental alleles in the hybrid. A) represents the global view using the linear model method and B) represents the method used by (McManus et al. 2010)

**Table 3.** Regulatory category assignments of genes using the linear model (limma) method.

| Category | Treatment | Barke | Igri | BCC131 | HOR1969 | FT11 | FT67 | FT279 | FT581 |
| --- | --- | --- | --- | --- | --- | --- | --- | --- | --- |
| Cis only | Control | 283 | 340 | 368 | 811 | 1,065 | 962 | 894 | 953 |
|  | Cold | 8 | 230 | 352 | 782 | 748 | 904 | 967 | 1,033 |
|  | Intersection | 8 | 172 | 289 | 617 | 641 | 754 | 749 | 789 |
| Trans only | Control | 0 | 0 | 3 | 3 | 1 | 3 | 2 | 4 |
|  | Cold | 0 | 0 | 1 | 1 | 3 | 0 | 1 | 1 |
|  | Intersection | 0 | 0 | 0 | 1 | 0 | 0 | 0 | 0 |
| Cis + trans | Control | 0 | 0 | 3 | 15 | 7 | 14 | 15 | 12 |
|  | Cold | 0 | 0 | 1 | 9 | 7 | 14 | 18 | 18 |
|  | Intersection | 0 | 0 | 1 | 7 | 3 | 9 | 12 | 6 |
| Cis x trans | Control | 0 | 1 | 3 | 1 | 5 | 0 | 3 | 9 |
|  | Cold | 0 | 1 | 2 | 0 | 1 | 0 | 3 | 4 |
|  | Intersection | 0 | 1 | 1 | 0 | 1 | 0 | 1 | 2 |
| Compensatory | Control | 0 | 3 | 35 | 22 | 29 | 17 | 28 | 29 |
|  | Cold | 0 | 1 | 39 | 25 | 29 | 16 | 29 | 30 |
|  | Intersection | 0 | 1 | 28 | 17 | 20 | 12 | 19 | 20 |
| Conserved | Control | 3,969 | 3,924 | 1,906 | 5,428 | 7,278 | 6,704 | 6,610 | 7,024 |
|  | Cold | 3,960 | 3,916 | 1,938 | 5,514 | 7,871 | 6,945 | 6,654 | 7,093 |
|  | Intersection | 3,738 | 3,713 | 1,771 | 5,165 | 7,047 | 6,408 | 6,241 | 6,582 |
| Ambiguous | Control | 674 | 366 | 271 | 570 | 933 | 890 | 730 | 909 |
|  | Cold | 958 | 486 | 356 | 519 | 659 | 711 | 610 | 761 |
|  | Intersection | 454 | 159 | 87 | 179 | 229 | 300 | 222 | 283 |
| Total | Control | 4,926 | 4,634 | 2,589 | 6,850 | 9,318 | 8,590 | 8,282 | 8,940 |
|  | Cold | 4,926 | 4,634 | 2,589 | 6,850 | 9,318 | 8,590 | 8,282 | 8,940 |

**Table 4.** Regulatory category assignments of genes using the binomial and Fisher’s exact test method of McManus *et al*. (2010).

| Category | Treatment | Barke | Igri | BCC131 | HOR1969 | FT11 | FT67 | FT279 | FT581 |
| --- | --- | --- | --- | --- | --- | --- | --- | --- | --- |
| Cis only | Control | 1,459 | 1,118 | 759 | 1,738 | 2,704 | 2,546 | 2,338 | 2,303 |
|  | Cold | 1,084 | 1,294 | 749 | 1,924 | 2,073 | 2,458 | 2,239 | 3,070 |
|  | Intersection | 744 | 502 | 413 | 983 | 1,041 | 1,457 | 1,134 | 1,565 |
| Trans only | Control | 287 | 448 | 139 | 1,174 | 1,120 | 1,116 | 1,075 | 731 |
|  | Cold | 274 | 462 | 241 | 851 | 863 | 821 | 1,164 | 692 |
|  | Intersection | 41 | 84 | 26 | 319 | 249 | 265 | 283 | 142 |
| Cis + trans | Control | 241 | 806 | 248 | 1,012 | 1,110 | 969 | 1,082 | 580 |
|  | Cold | 158 | 472 | 228 | 763 | 1,262 | 586 | 1,430 | 741 |
|  | Intersection | 70 | 292 | 101 | 416 | 459 | 308 | 618 | 251 |
| Cis x trans | Control | 77 | 173 | 107 | 296 | 285 | 246 | 298 | 235 |
|  | Cold | 48 | 143 | 117 | 241 | 311 | 176 | 344 | 189 |
|  | Intersection | 14 | 29 | 39 | 94 | 100 | 69 | 100 | 50 |
| Compensatory | Control | 89 | 291 | 200 | 212 | 272 | 193 | 247 | 313 |
|  | Cold | 40 | 168 | 90 | 208 | 467 | 154 | 219 | 191 |
|  | Intersection | 5 | 24 | 12 | 34 | 44 | 23 | 25 | 22 |
| Conserved | Control | 1,173 | 909 | 653 | 1,170 | 1,995 | 1,578 | 1,617 | 2620 |
|  | Cold | 968 | 925 | 623 | 1,417 | 2,458 | 2,127 | 1,280 | 1,932 |
|  | Intersection | 469 | 354 | 319 | 526 | 1,068 | 889 | 556 | 1,058 |
| Ambiguous | Control | 1,600 | 889 | 483 | 1,248 | 1,832 | 1,942 | 1,625 | 2,158 |
|  | Cold | 2,354 | 1,170 | 541 | 1,446 | 1,884 | 2,268 | 1,606 | 2,125 |
|  | Intersection | 992 | 330 | 144 | 408 | 541 | 725 | 506 | 670 |
| Total | Control | 4,926 | 4,634 | 2,589 | 6,850 | 9,318 | 8,590 | 8,282 | 8,940 |
|  | Cold | 4,926 | 4,634 | 2,589 | 6,850 | 9,318 | 8,590 | 8,282 | 8,940 |

The numbers of genes in each regulatory category are roughly similar for control samples and those in response to environmental stress, but since these tests were conducted independently, we wanted to know how similar these lists are. To answer this question, we found the intersection of gene lists for each comparison (Table 3-4). The results show that regulatory category assignments are robust to environmental stress, especially for genes with conserved regulation. On average, 94% (∼90-96%) of genes in this category are present in both treatments for all crosses. Since it appears that results for category assignments are similar between treatments, we wanted to know what if we could detect more *trans* effects by considering control and cold treatments together to gain additional replicates, in order to gain statistical power within the linear model. The results (Table 5, Figure S8) are similar to when each treatment was analyzed separately (Table 3). A moderate increase in the number of *trans*, *cis* + *trans* and *cis* × *trans* effects were observed, but not to the same extent as found by (McManus et al. 2010).

**Table 5.**
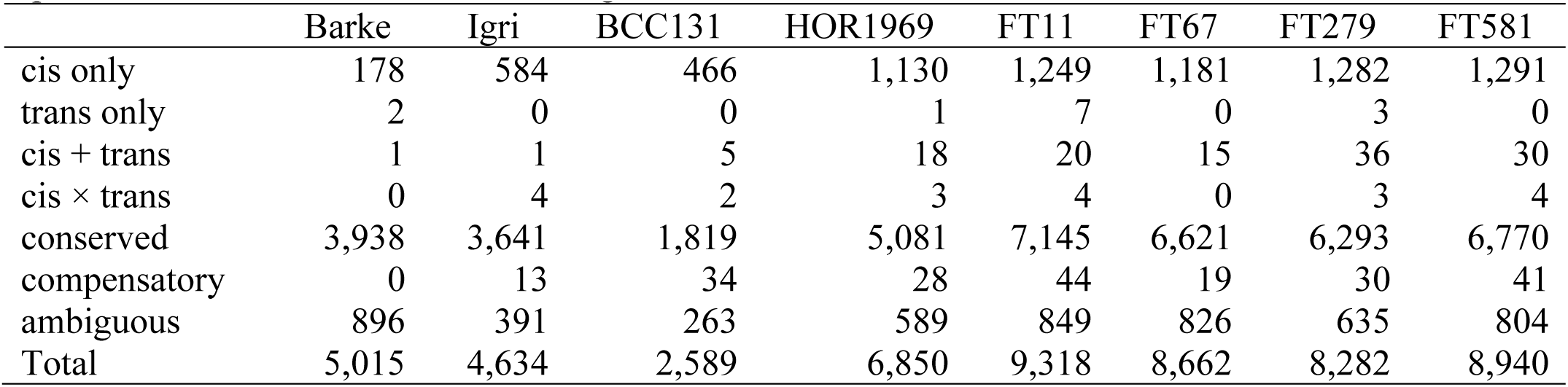
Regulatory category assignments for each cross when treatments were not considered separately and instead grouped as additional replicates. The linear model was used to generate these results..

### Expression of known cold responsive genes

In addition to looking at general expression patterns, we are also interested in the expression of known cold-responsive genes. Therefore, we looked into the expression patterns of these genes including *Vernalization1* (*VRN1*) and *Cold-Regulated 14B* (*COR14B*). The expression of both *VRN1* (HORVU5Hr1G095630) and *COR14B* (HORVU2Hr1G099830) matched our expectations. Morex and Barke (spring types) have higher expression levels of *VRN1* than Igri (a winter type), both landraces, as well as all wild accessions (Figure 5A). Expression of *VRN1* is maintained at low levels in wild and winter barleys until it has endured a prolonged period of cold exposure, or vernalization. This vernalization requirement is evolutionarily advantageous because flowering will only occur when prevailing environmental conditions are favorable.

**Figure 5.**
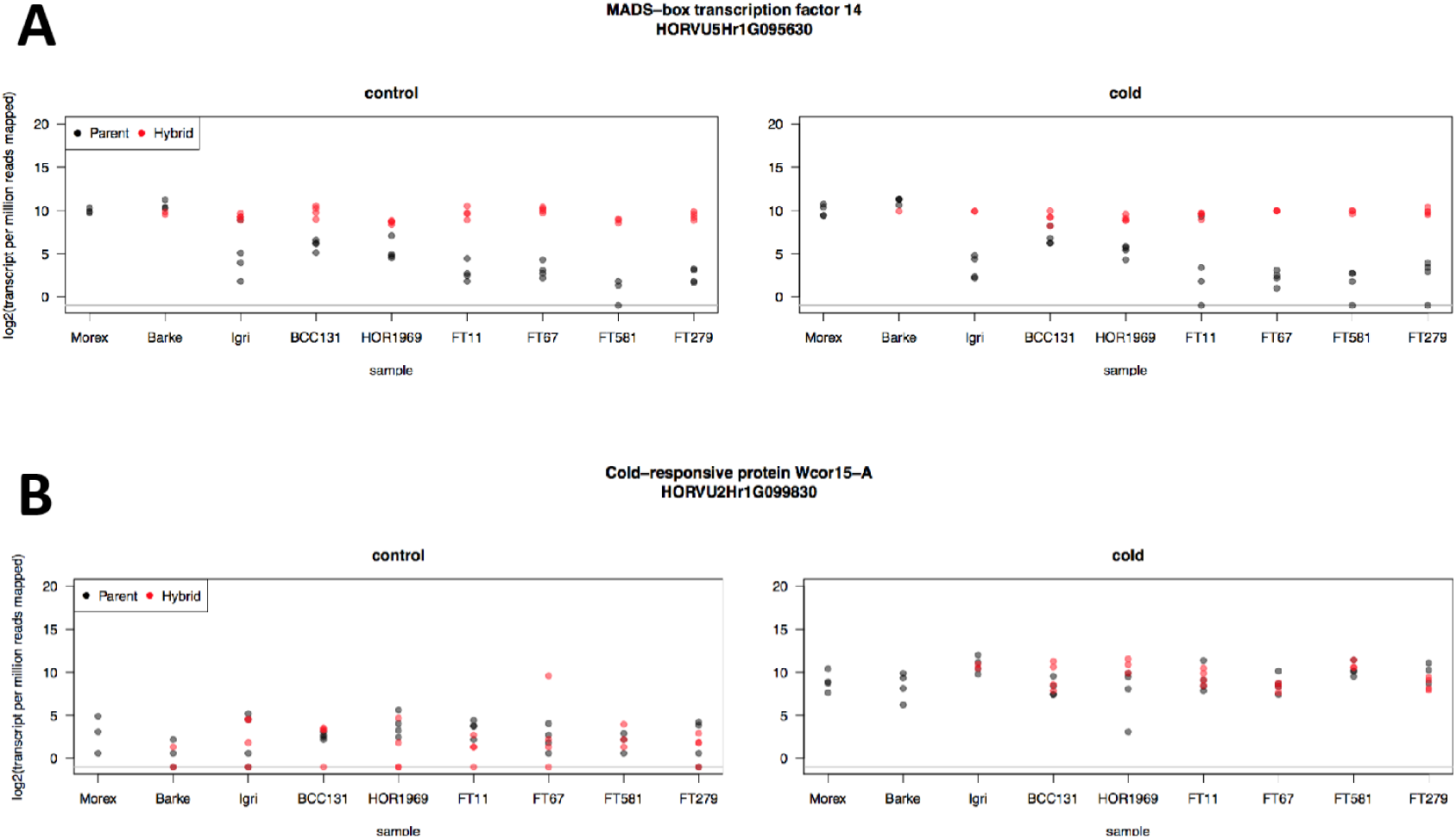
Expression (log_2_-transformed transcripts per million reads mapped) values for parents (black) and hybrids (red) from each sample: A) *VRN1* (HORVU5H1rG095630) is expressed at higher levels in spring types (which do not require vernalization) than the winter type, landraces and wild barleys. The hybrids show this same higher level of *VRN1* expression, indicating that the Morex allele is dominant. B) *COR14B* (HORVU2Hr1G099830) shows a response to chilling in the cold-treated samples, also as expected.

Hybrids have *VRN1* expression levels that match those of the spring types, demonstrating that the loss of a vernalization requirement is dominant. The expression profiles are similar for both control and cold treatments, which is also expected since *VRN1* expression only increases after several weeks of exposure to cold temperatures and our samples were only exposed to cold for three hours. The expression of *VRN1* in the hybrid confirms that the hybrid shows the correct inheritance patterns. Therefore, we believe that our other results are reliable. The cold-responsive gene *COR14B*, however, shows a clear increase in response to cold treatment (Figure 5B). This is also expected, since the plant response to chilling occurs rapidly (Cattivelli and Bartels 1989).

### Dominant vs. additive inheritance

In addition to regulatory categories discussed earlier, we are also interested in examining the mode of inheritance of genes in our dataset. Many genes exhibit Mendelian inheritance (dominance vs recessive). However, many other genes exhibit quantitative or additive inheritance. Still other inheritance modes (heterosis) also exist. Heterosis, also known as hybrid vigor, occurs when expression of a gene is outside of the range of the parental values (i.e., overdominance). We were interested in exploring the distribution of these inheritance modes in our data. The summary of the modes of inheritance is reported in Table 6 for control samples and Table 7 for cold samples. Numbers are unavailable for Morex × Barke under the cold treatment because of a lack of replicates for cold Barke hybrid samples. Relatively few genes show heterotic effects (overdominance) under both control and cold conditions. For most crosses, these categories represent less than 1% of differentially expressed genes. Under no circumstance did heterosis affect more than 2% of differentially expressed genes. Approximately one third of all differentially expressed genes have additive effects under both conditions. (25.5%-37.8% control and 30.3%-38.0% cold). Genes showing dominance together represent about another third of differentially expressed genes. In nearly every cross, more Morex alleles are dominantly expressed than the paternal allele. This could be an effect of Morex being the maternal allele, but it could also reflect a tendency of domesticated alleles to be more highly expressed than wild alleles. The one case where the paternal allele has more dominantly expressed alleles than Morex involved Igri, a winter cultivar, under control conditions. Otherwise, the trend seems to be that the numbers of dominant genes are more equally distributed between the two parents for cultivars (Barke and Igri) and landraces (BCC131 and HOR1969) than for the wild accessions (FT11, FT67, FT279 and FT581). Another quarter to one third (27.4%-36.2% control and 24.5%-34.3% cold) of all differentially expressed genes were placed into the ambiguous category and a handful of others did not fit into any of the other categories. It is difficult to speculate which category these genes truly belong to. We might assume that they would fall into one of the main three categories (Additive, Morex dominant or paternal allele dominant) in a proportional manner, but we cannot state this with certainty.

**Table 6.**
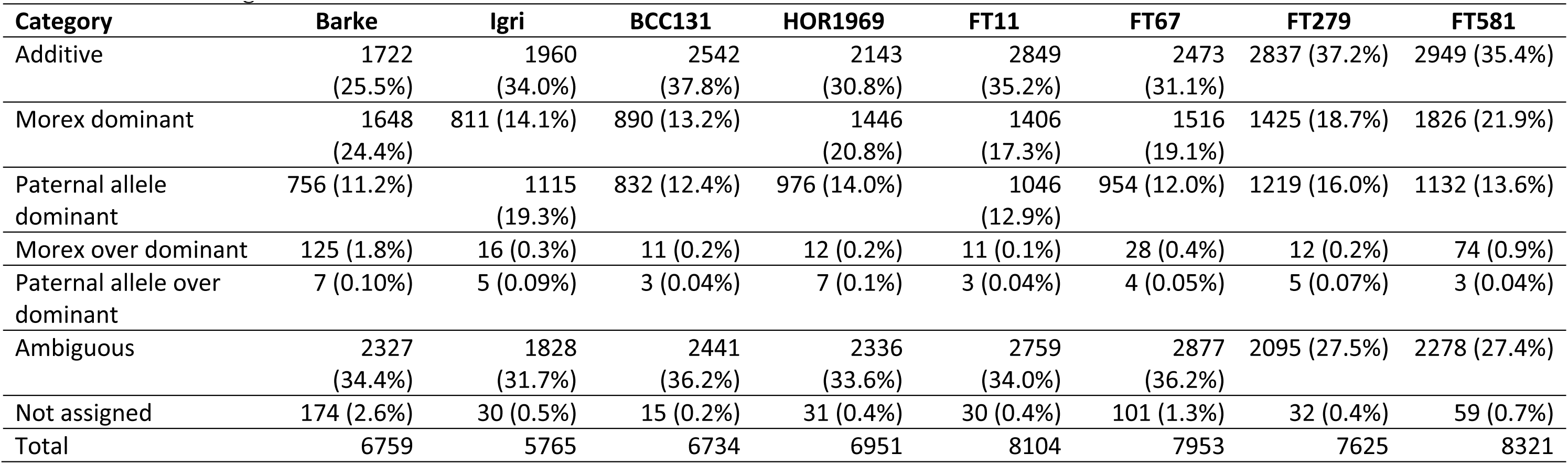
Mode of inheritance assignment counts for each cross under control conditions. Percentage values may not add up to exactly 100% due to rounding.

| Category | Barke | Igri | BCC131 | HOR1969 | FT11 | FT67 | FT279 | FT581 |
| --- | --- | --- | --- | --- | --- | --- | --- | --- |
| Additive | 1722<br>(25.5%) | 1960<br>(34.0%) | 2542<br>(37.8%) | 2143<br>(30.8%) | 2849<br>(35.2%) | 2473<br>(31.1%) | 2837 (37.2%) | 2949 (35.4%) |
| Morex dominant | 1648<br>(24.4%) | 811 (14.1%) | 890 (13.2%) | 1446<br>(20.8%) | 1406<br>(17.3%) | 1516<br>(19.1%) | 1425 (18.7%) | 1826 (21.9%) |
| Paternal allele dominant | 756 (11.2%) | 1115<br>(19.3%) | 832 (12.4%) | 976 (14.0%) | 1046<br>(12.9%) | 954 (12.0%) | 1219 (16.0%) | 1132 (13.6%) |
| Morex over dominant | 125 (1.8%) | 16 (0.3%) | 11 (0.2%) | 12 (0.2%) | 11 (0.1%) | 28 (0.4%) | 12 (0.2%) | 74 (0.9%) |
| Paternal allele over dominant | 7 (0.10%) | 5 (0.09%) | 3 (0.04%) | 7 (0.1%) | 3 (0.04%) | 4 (0.05%) | 5 (0.07%) | 3 (0.04%) |
| Ambiguous | 2327<br>(34.4%) | 1828<br>(31.7%) | 2441<br>(36.2%) | 2336<br>(33.6%) | 2759<br>(34.0%) | 2877<br>(36.2%) | 2095 (27.5%) | 2278 (27.4%) |
| Not assigned | 174 (2.6%) | 30 (0.5%) | 15 (0.2%) | 31 (0.4%) | 30 (0.4%) | 101 (1.3%) | 32 (0.4%) | 59 (0.7%) |
| Total | 6759 | 5765 | 6734 | 6951 | 8104 | 7953 | 7625 | 8321 |

**Table 7.**
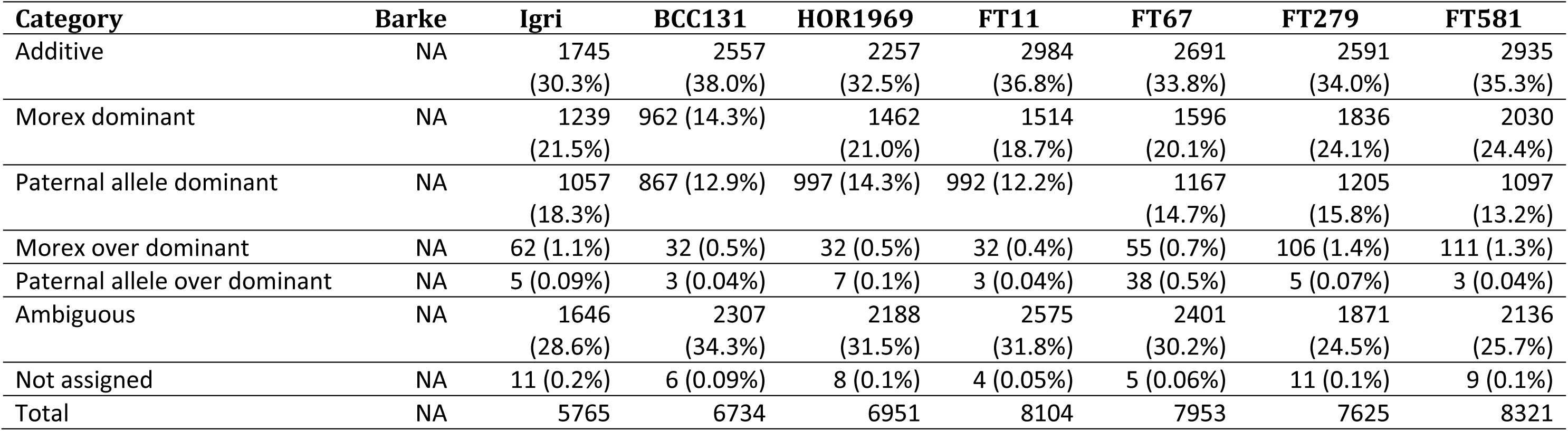
Mode of inheritance assignment counts for each cross under cold (4°C) conditions. Values for Barke are unavailable because of a lack of replicates from hybrid in the cold treatment. Percentage values may not add up to exactly 100% due to rounding.

## Discussion

We are interested in understanding the effect of domestication on patterns of gene expression and regulatory variation in barley. To accomplish this, we combined the use of ASE on a small panel of wild and domesticated barleys and their F_1_ hybrids with a cold stress treatment according to established methods (Cowles et al. 2002; Cubillos et al. 2014; Lemmon et al. 2014). Several lines of evidence indicate that the approach worked and our results are reliable. First, samples cluster according to generation (Figure 2A), accession (Figure 2B) and treatment (Figure 2C). The expression profiles of cold-responsive genes such as *VRN1* and *COR14B* also behave as expected (Figures 5A-5B). Of 39,734 high-confidence genes in the barley genome, we were able to quantify ASE for between 2,589 (BCC131) and 8,940 (FT581) genes (Table 2). We cannot measure ASE for genes that lack SNPs because it is impossible to unambiguously assign such transcripts to a parental allele without at least one SNP to verify the allele of origin. Other genes may not be expressed at sufficient levels to have statistical power for ASE. Based on previous studies (McManus et al. 2010; Lemmon et al. 2014), we expected to find a similar number of genes regulated in *cis* and *trans*; however, we found almost a complete absence of genes regulated in *trans*. The increased expression of cold response genes (*COR14B*, Figure 5B) after cold treatment suggest that the cold treatment induced transcription factors (TFs) to elicit a response to cold. Since TFs act in *trans*, some *trans* effects are expected; however, a small number of TFs may be more plausible than hundreds or thousands of *trans*-acting genes observed in earlier studies, to minimize pleiotropic effects (West et al. 2007). In general, genes with *trans* effects may not cause pleiotropic effects if they do not disrupt highly connected nodes in a network (Jeong et al. 2001; Fraser et al. 2002). Further, TFs do not necessarily cause large pleiotropic effects. Work in *C. elegans* shows that mutations in the Ras signaling pathway that activate multiple TFs are more deleterious than mutations affecting only TFs (Kayne and Sternberg 1995). In our present study, the genes regulated in *trans* according to the linear model do not appear to have any great significance. The expression levels of these genes are low and are plagued with missing data (e.g., some of the genes are expressed in one genotype, but not another) and annotations are ambiguous. It is also possible that the parameters of our analysis are too strict, resulting in false negatives; however, other studies have likely suffered from false positives. Clearly, a method is needed that rejects *trans* effects that are truly absent, but accepts real *trans* effects.

Evidence for regulatory changes in response to environmental stress is absent from our data, in agreement with Cubillos et al. (2014). However, we cannot rule out that the use of a different environmental stress (high temperature, drought or salinity) could induce a more variable response. Cubillos et al. (2014) also found that roughly half of the genes in their samples had compensatory effects, meaning that *cis* and *trans* effects have opposite effects. In contrast, in we found that half of our genes had conserved effects. In addition, Cubillos et al. (2014) observed an increase in the number of genes with *trans* effects that resulted in a change in direction in response to the environment, rather than a change in magnitude, compared to genes with *cis* effects. We were not able to make such a comparison, since genes with *trans* effects are virtually absent in our dataset.

Wittkopp et al. (2008) found a greater amount of *cis* regulatory expression differences between species rather than within species, which could also explain why *trans* effects were more pronounced in studies that examined expression differences between *Drosophila* species (McManus et al. 2010). However, Osada et al. (2017) also noted large variances in their samples; therefore, our hypothesis that differences observed for *trans* regulation are likely to be false positives as a result of statistical artifacts seems to be plausible.

The observation of a greater number of *cis*- compared with *trans*-acting factors has important implications for the use of crop wild relatives in plant breeding. Insights into gene regulation in barley such as this will help to exploit wild genetic resources in elite germplasm (Schmalenbach et al. 2009). In nature, it appears that *cis* effects preferentially accumulate, likely due to fewer pleiotropic effects compared with *trans* effects (Prud’homme et al. 2007). Similarly, in plant breeding, genetic background is known to influence the expression of genes due to epistatic interactions (Kroymann and Mitchell-Olds 2005; Blanc et al. 2006). For novel quantitative trait loci (QTL) introgressed into elite germplasm to be useful, the beneficial trait must be expressed in the elite background. Genes regulated in *cis* will be more likely to be expressed at the same level in a novel background as in their native background when their regulatory sequence is co-inherited due to linkage, whereas co-inheritance of *trans* regulators will occur less frequently due to independent segregation. Introgression of a gene as well as its *trans* regulator would be complicated enough, but could also have deleterious effects in the new genetic background if the *trans* regulator epistatically affects the expression of off-target genes. The recipient background may also regulate the introgression through *trans* regulators. One way to study this experimentally is to use near-isogenic lines (NILs) that contain as many of the total possible genes in small introgressions throughout the genome. Guerrero et al. (2016) conducted such an experiment in tomato. They showed that introgressed genes tend to be down regulated while native (non-introgressed) regions tend to be up regulated. The authors concluded that c*is*- and *trans*-regulation have roughly equal contributions to expression divergence.

The *cis* regulatory regions of genes can be large, extending for thousands of kilobases such as the case with *Teosinte Branched 1* (*tb1*) in maize, which has at least one regulator from 58-69 kb upstream from the 5’ start site (Clark et al. 2006). Therefore, it is possible that recombination may occur between a *cis*- regulatory sequence and the gene it controls. However, *cis* regulatory regions are not well defined. This possibility highlights one limitation of the applications of our study. Due to our experimental design, we can only infer the presence and relative contribution of *cis*- or *trans*-acting regulation, but we cannot map these regulators; therefore, we do not know the genomic position of these regulators. An experimental approach known as expression quantitative trait loci (eQTL) mapping allows gene expression to be mapped as quantitative traits in experimental populations or by association genetics. These studies allow for mapping of regulatory elements; however, it is still not always clear at what distance threshold an eQTL would be acting in *cis* or in *trans*, since these distance thresholds are often arbitrary (Lagarrigue et al. 2013). In addition, eQTL are more properly referred to as local or distant, rather than *cis* or *trans* (Lagarrigue et al. 2013). These studies are also more difficult and expensive because they require a large mapping population to be both genotyped and assayed for genome-wide expression values.

Alignment bias due to polymorphism or structural variants is a well-documented problem with ASE studies (Degner et al. 2009; Stevenson et al. 2013) and our dataset is no exception due to the use of a single genotype (Morex) as a reference. In part due to these limitations, a single reference genotype is no longer considered to be sufficient to capture the full diversity present in a given species. The concept of the pan-genome posits that any species has a set of genes present in all accessions (the core genome), genes that are present in some, but not all accessions (the dispensable genome) and lineage-specific genes that are only present in a single accession. In this context, additional reference genomes are needed. Other barley genotypes, such as Barke and FT11 are not available at present. There is a barley pan-genomic project underway, at which point these genotypes and others will be available. Please see Monat et al. (2018) for a summary of this topic. For now, it is necessary to interpret our results with caution. When the genomes of these other accessions do become available, it will become possible to re-analyze these data to measure the impact of the reference bias.

The availability of additional reference genomes will also allow for re-analysis of these data with a Bayesian approach that allows for direct comparison of environmental effects (León-Novelo et al. 2018). Additional reference genomes are necessary because the method incorporates the number of RNA reads which align equally well to both parental genomes.

## Materials and Methods

### Growth conditions

Plants were grown in a growth chamber with a 12 h photoperiod with temperatures of 22°C and 18°C during light and dark periods, respectively. After one week of growth, when the first leaf of each accession was fully expanded, half of the plants were moved to a cold room at 4°C for 3 h. The response to chilling occurs rapidly in barley (Cattivelli and Bartels 1989), so this short cold treatment is sufficient to induce a physiological response. After the 3 h cold treatment, the first leaf of each individual from both groups was harvested and immediately frozen in liquid nitrogen before being moved to storage at −80°C. Each cold treatment (11:00) and tissue harvest (14:00) was conducted at the same time of the day for each replicate to avoid confounding factors associated with circadian rhythm. The experimental design is shown in Figure 1. The experiment was replicated four times. For accessions that either failed to germinate or grew poorly, a fifth attempt was made to obtain additional replicates. As a result, most samples were replicated four times. A few samples have only three replicates: FT67 hybrid cold, FT581 parent control, both FT581 hybrid control and cold, and Morex parent control. Two samples have only two replicates: Barke hybrid control and Igri hybrid cold. One sample, Barke hybrid cold, was not able to be replicated despite repeated efforts to get more data.

### RNA extraction, sequencing and data analysis

Frozen leaf tissue (−80°C) was homogenized by grinding to a fine powder in 1.5 ml tubes with metal beads two times for 30 s each (1 min total) at 30 Hz using a mixer mill (Retsch GmbH, Haan, Germany). Tubes containing the samples were submerged in liquid nitrogen between grinding to ensure samples did not thaw during the process. Once all samples were ground, RNA was extracted using RNeasy^®^ mini kits (Qiagen) according to manufacturer’s instructions. To remove any DNA contamination, samples were treated with Ambion^TM^ DNase (ThermoFisher Scientific) according to manufacturer’s instructions. RNA quality and integrity were checked with an Agilent 2100 Bioanalyzer (Agilent Technologies) and a Qubit^TM^ 2.0 fluorometer (ThermoFisher Scientific), respectively.

Where possible, three individuals of each parent or hybrid were planted for each replicated treatment. The healthiest plant (e.g., not yellow or stunted) was selected for harvesting. After RNA extraction was carried out according to the methods described above, high quality RNA (mass ≥ 1 μg, volume ≥ 20 μL, concentration ≥ 50 ng/μL, RIN ≥ 6.3, and 260/280 and 260/230 ≥ 2.0) samples were submitted for sequencing.

In total, 123 NEB Next^®^ Ultra^TM^ RNA libraries with an average insert size of 250-300 bp were sequenced (paired-end, 2× 150 cycles) on an Illumina HiSeq 2500 machine. RNA sequencing was done by Novogene while exome capture sequencing was performed at the IPK sequencing center. RNA-seq data were quantified using both the pseudoalignment software kallisto v. 0.43.0 (Bray et al. 2016) and HISAT2 (Kim et al. 2015). The abundance files from kallisto and HISAT2 were separately loaded into the R statistical environment (R Core Team 2012) for further analysis. Gene abundance estimates from kallisto were normalized using edgeR and limma (Robinson et al. 2010; Ritchie et al. 2015) and the voom transformation (Law et al. 2014) was applied to account for the mean-variance relationship of RNA-seq data. These data were used to calculate the variance using the matrixStats package (Bengtsson 2016). The 1000 genes with the highest variance were used for PCA. Kallisto was used to find overall expression patterns while HISAT2 was used for allele-specific expression. All raw RNA sequence data are available from the European Nucleotide Archive (ENA) under accession numbers PRJEB29972. Accession numbers for individual samples are provided in Table S1.

### DNA extraction and exome capture

In order to select high-confidence variants for allele-specific expression analysis using a genomic control, an exome capture assay was applied for the eight hybrid genotypes (Mascher et al. 2013). Exome capture data for the parental genotypes may be found in Russell et al. (2016). Genotype matricies for SNPs (https://doi.org/10.5447/IPK/2016/4) and indels (https://doi.org/10.5447/IPK/2016/5) for parental accessions are available through e!DAL (Arend et al. 2014). The raw sequence data for these parents were deposited into the ENA and the accession codes are available in Supplementary Table 1 of Russell et al. (2016). For the present study, hybrid DNA was extracted using a DNeasy^®^ kit (Qiagen). DNA concentrations were measured using a Qubit^TM^ 2.0 fluorometer (ThermoFisher Scientific) and all samples were above 20 ng μl^-1^. DNA integrity was verified using a 0.7% agarose gel, which showed that DNA from each sample was intact. Sequencing was performed using an Illumina Hiseq 2500 machine (2 x 100 bp, insert size = 320 bp). Captured reads were mapped against the BAC-based Morex reference sequence (Mascher et al. 2017) with BWA-MEM (Li et al. 2013). Coverage was determined using the depth command from SAMtools (Li 2011) using only properly paired reads. Mapping statistics are available in Figure S3.

### Allele-specific transcript quantification and normalization

The R package limma (Ritchie et al. 2015) was used for the analysis of ASE using a linear model approach. Briefly, allele-specific counts were converted into a matrix and rounded to the nearest integer. Counts were then normalized using edgeR (Robinson et al. 2010) to account for differences in total read count between samples and stored in a differential expression list. A design matrix was created using each combination of generation × accession × treatment as a single factor. The voom transformation was applied to the count matrix to account for the mean-variance relationship of RNA-seq data. The linear model was created by fitting the voom-transformed (Law et al. 2014) count matrix to the design matrix. Differentially expressed genes were identified using the contrasts specified in the contrast matrix. For example, the expression level of each individual parent was contrasted to Morex to decide whether the parents were different from each other. Subsequently, the parental alleles within the hybrid were compared to each other to decide if their expression was different from each other.

These allele-specific counts were also used as input for differential expression analysis. We used the differential expression analysis to assign the mode of inheritance for differentially expressed genes, which is described below.

### Assignment of regulatory categories

To find variants between samples, SNPs were called from sorted and indexed binary alignment map (BAM) files originating from exome capture and RNA-seq samples. The BAM files were sorted and indexed using Novosort (http://www.novocraft.com/products/novosort). Results were imported into R for further analysis. The SNP matrix was assigned a Digital Object Identifier (https://doi.ipk-gatersleben.de/DOI/43e62feb-1fd8-42a0-af62-f5e1a872b61c/4c61bc40-da8f-4fd5-9486-dd0ce183c205/2/1847940088) and registered with e!DAL (Arend et al. 2014). Raw DNA sequence data are available through the ENA under accession number PRJEB29973.

Allele-specific counts were derived from SNPs in the RNA-seq data that were corroborated by a genomic control. First, informative SNPs were detected in the exome capture data. SNPs were considered informative in a specific cross if they the parents carried different alleles in homozygous state. In addition, the successful genotype calls in the hybrid exome capture data were required. Then, we determined how many reads supported the reference allele or the alternate allele in RNAseq data for parents hybrid and calculate the DV/DP ratio (depth of the variant allele vs. total read depth). Information for multiple SNPs were combined at the gene level by merging the SNP information with gene information in the R statistical environment and summing up DP and DV values for all SNPs in a gene. Low DV/DP ratios indicate that more reads originated from the reference (maternal = Morex) allele while a high DV/DP ratio indicate more reads originated from the alternate (paternal) allele. A DV/DP ratio of 0.5 means that both alleles are expressed equally. Genes with less than 50 reads across all samples were filtered out before further analysis. The design matrix was created by considering each combination of accession, generation and treatment as a single factor. The linear model created by fitting the model specified in the design matrix to the voom-transformed (Law et al. 2014) count matrix. Genes may be assigned to one of seven regulatory categories described by McManus et al. (2010). Genes with significant (FDR adjusted p-value ≤ 0.01 using Benjamini-Hochberg procedure) expression differences between parents and parental allele expression levels matching that of their respective parent in the hybrid were assigned to the *cis only* category (Figure S1A). In contrast, genes with significant expression differences between parents, but not between parental alleles in the hybrid were assigned to the *trans only* category (Figure S1B). Figures S1C and S1D show the expectations for cis + trans and *cis × trans* categories, respectively. Full descriptions of regulatory categories may be found in McManus et al. (2010).

### Dominant vs. additive inheritance

We used our gene expression dataset to find whether genes were inherited in a dominant or an additive manner. We use the classifications given by Albert et al. (2018) to make assignments. First, we used the subset of differentially expressed genes from each cross as described above. Genes were assigned as Morex dominant if the expression of the gene in the hybrid was greater than in the low parent and matching the expression of Morex. Genes were called recessive when the expression in the hybrid was lower than Morex and matched that of the low parent. We renamed these as “paternal allele dominant” in the final tables. Additive genes were those genes which had intermediate expression values between the two parental alleles. Genes which had higher expression values than both parents and Morex was the high parent were placed into the Morex overdominant category. Genes which had higher expression values than both parents and the paternal parent was the high parent were assigned to the “paternal allele overdominant” category. For the genes which remained unclassified, we used log_2_ FC expression values below 1 and greater than −1 for each contrast to assign these genes to the “ambiguous” classification. Even after this step, some genes remained unassigned. We report these genes as “not assigned”. The number of genes in each category was small, but for two crosses (Barke and FT67) the number of unassigned genes was relatively high at 174 and 101, respectively.

## Acknowledgements

We are grateful to Nils Stein for provision of seed material and to Manuela Knauft, Susanne König, Ines Walde and Mary Ziems for technical assistance. We also thank Andreas Czihal and Dominic Knoch for advice regarding RNA extractions and Anne Fiebig for uploading data to the ENA. We greatly appreciate the financial support from the German Research foundation (Deutsche Forschungsgemeinschaft, DFG) to Martin Mascher (Grant ID: MA6611/2).

## Supporting Information

**Figure S1.**
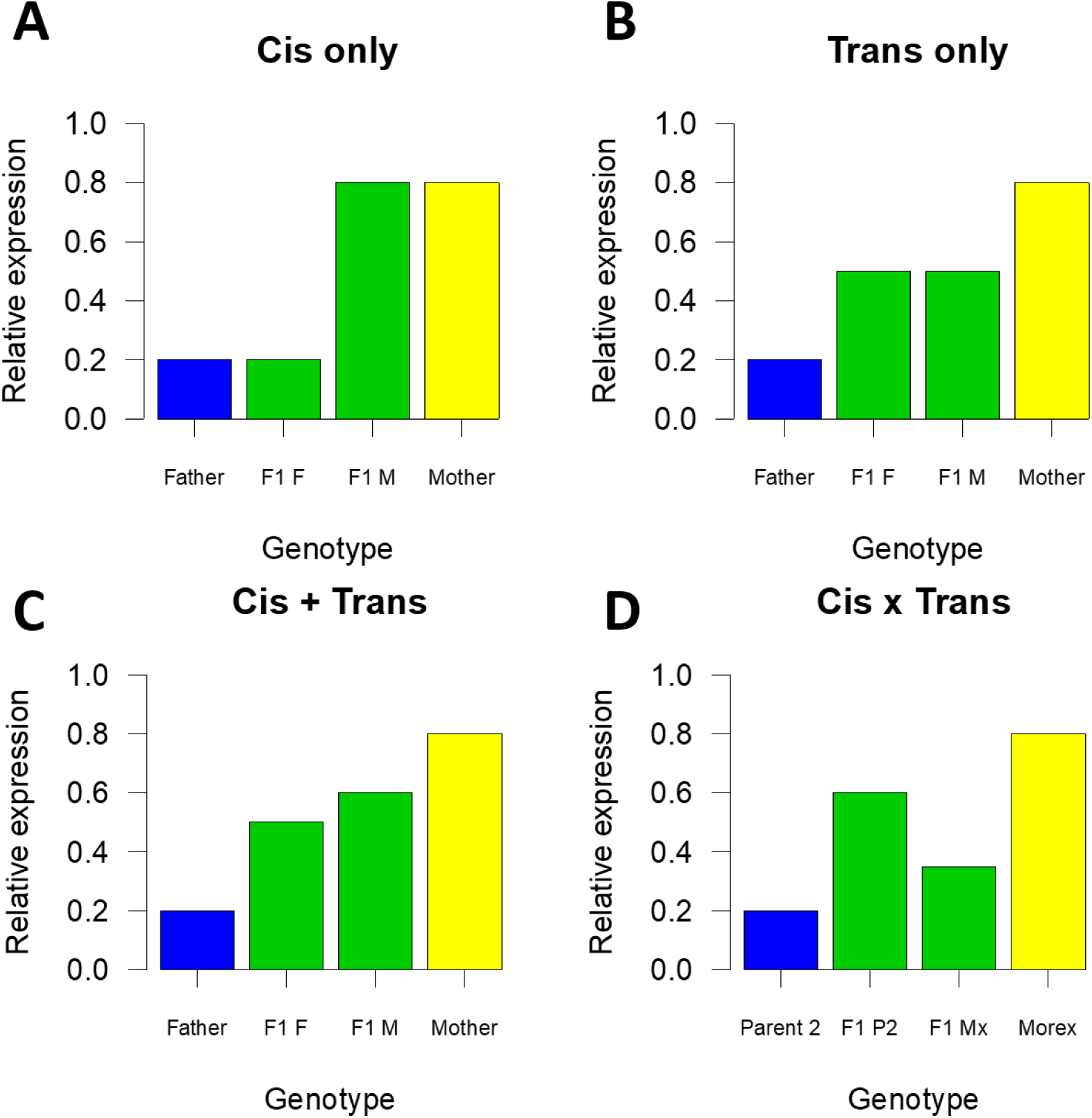
Expected relative expression levels for A) *Cis* only effects, B) *Trans* only effects, C) *Cis + Trans* and D) *Cis × Trans*. Bar plots for the father (blue) and mother (yellow) are the result of the combined effects of both alleles in the respective accession. Two green bars (middle) each represent a single allele in the hybrid individual. F1 F is the hybrid allele derived from the father while F1 M is the hybrid allele derived from the mother.

**Figure S2.**
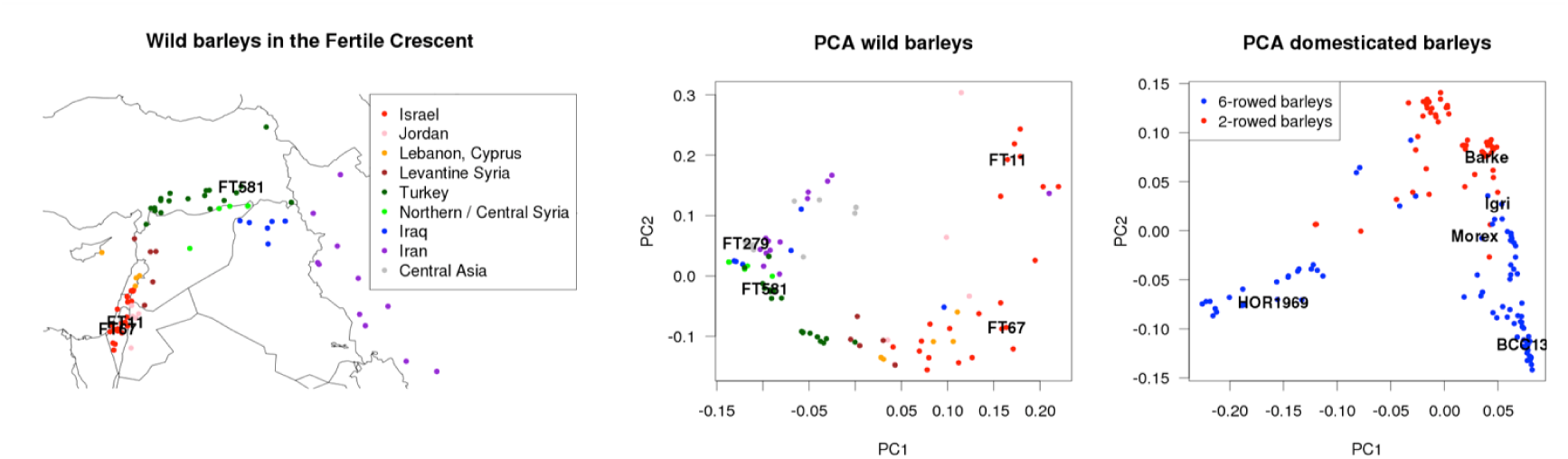
Geographical distribution of wild barleys used in this study (except FT279 from Afghanistan, which is not in the frame); and Principal component analysis based on exome capture data from (Russell et al. 2016) that was the basis of selection of parents for use in this study.

**Figure S3.**
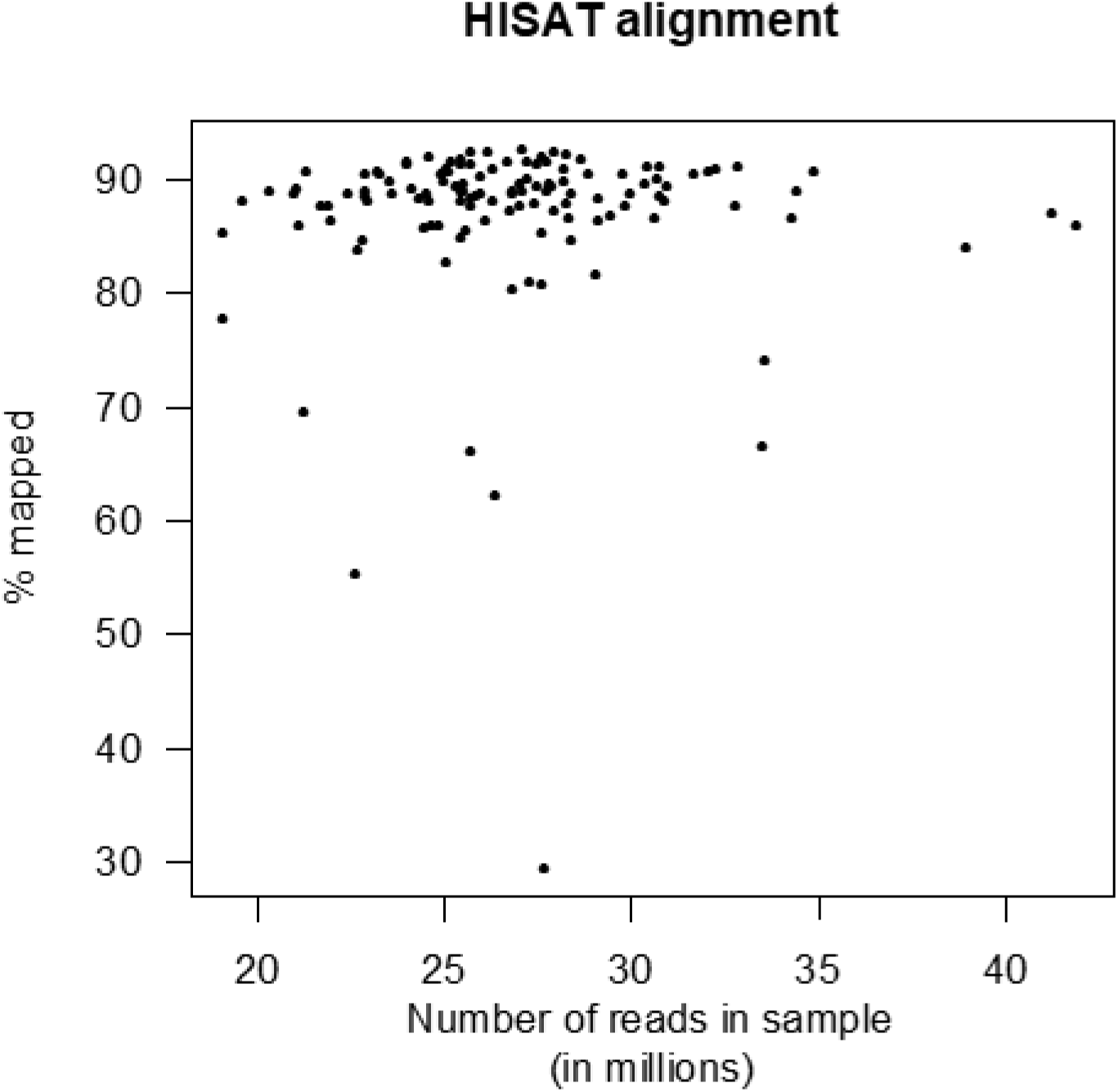
HISAT mapping rate.

**Figure S4.**
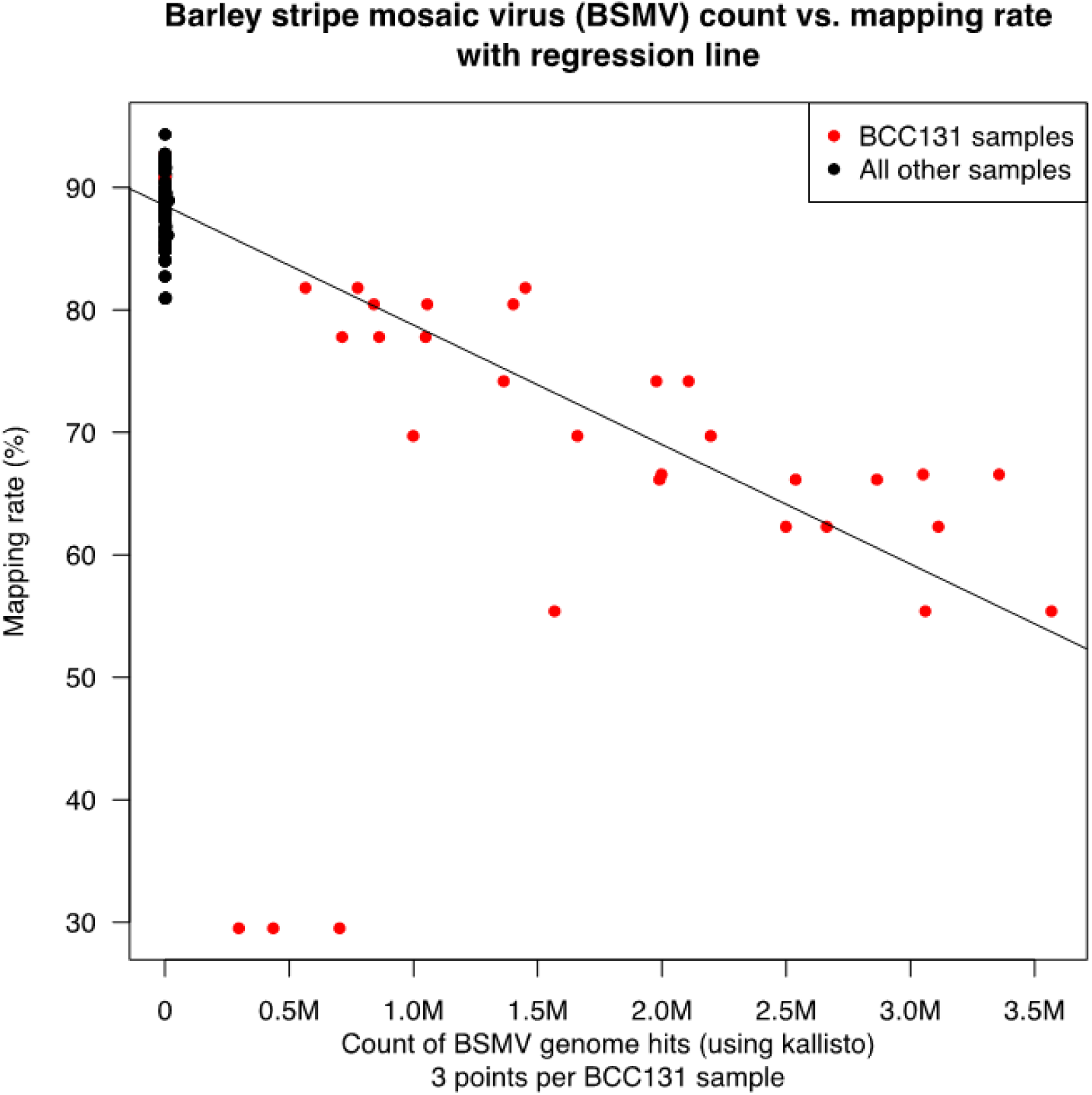
Barley stripe mosaic virus (BSMV) kallisto vs. HISAT mapping rate.

**Figure S5.**
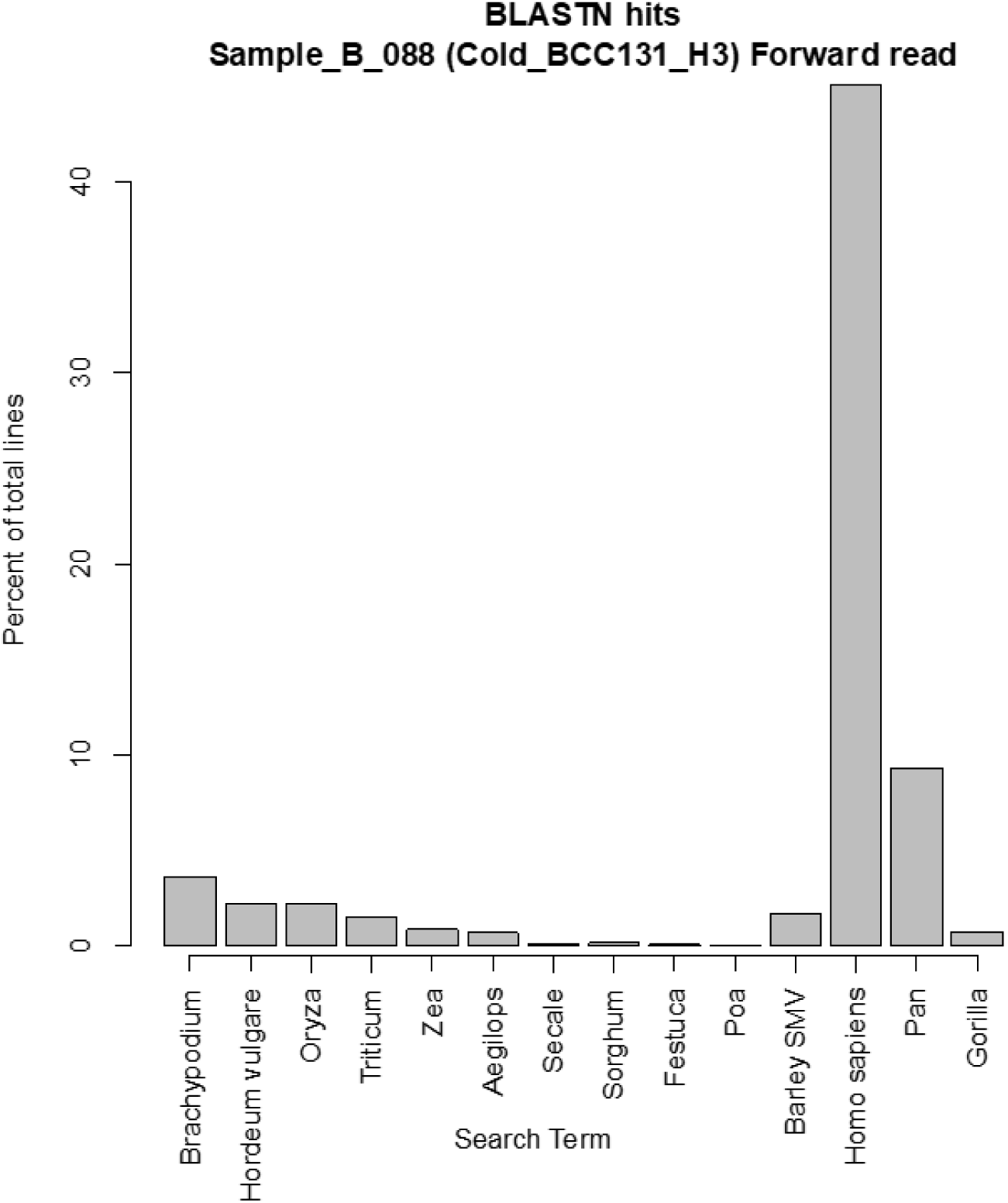
Basic Local Alignment Search Tool (BLAST) results for the forward read of Sample_B_088 (Cold_BCC131_H3).

**Figure S6.**
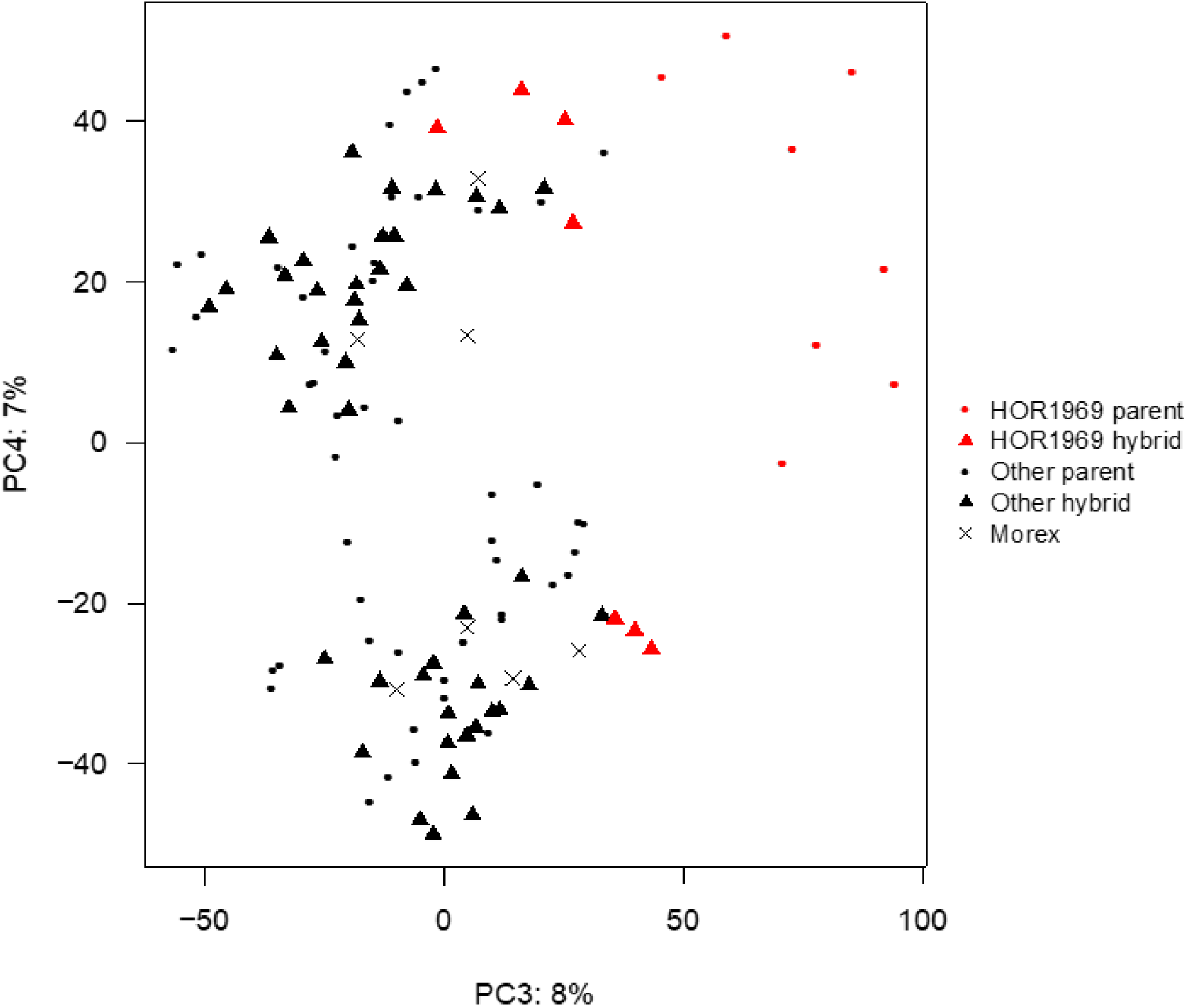
PCA plot of PC3-4. The plot is identical to the one presented in Figure 2C except that HOR1969 samples are colored in red and all other samples are colored in black. Sample shapes designate generation. Circles are parental samples while hybrids are triangles. Morex is indicated with a “×” symbol.

**Figure S7.**
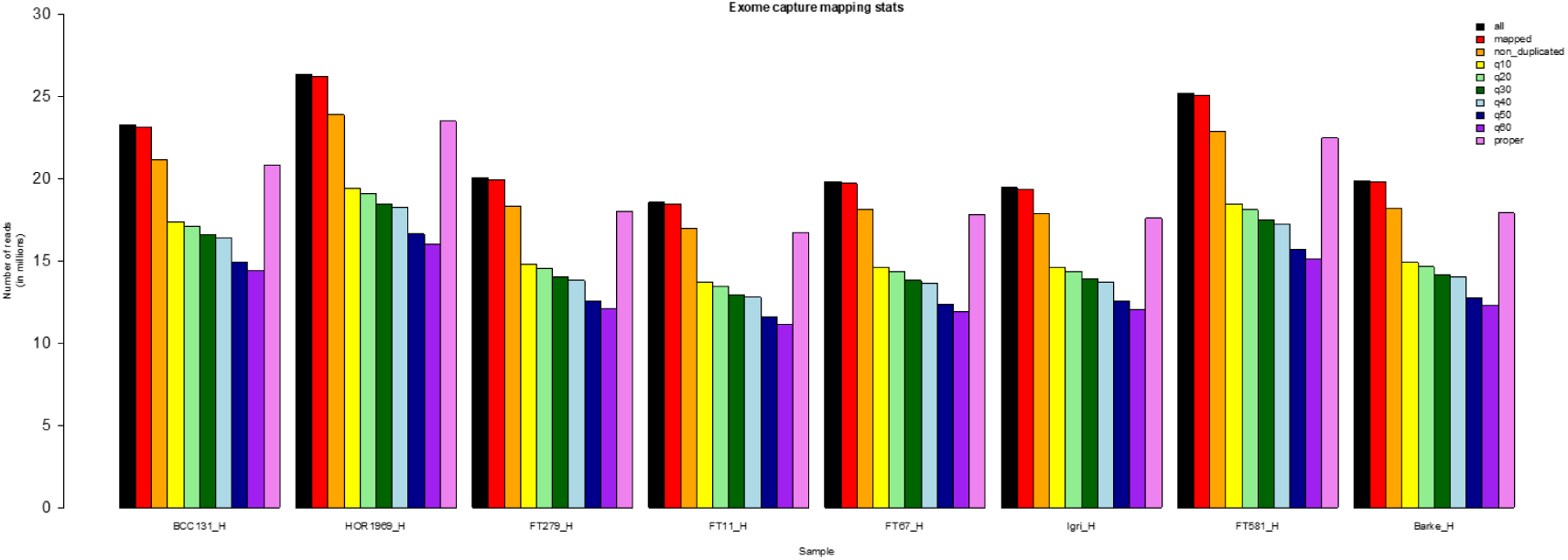
Exome capture mapping statistics for the eight hybrids used in this study.

**Figure S8.**
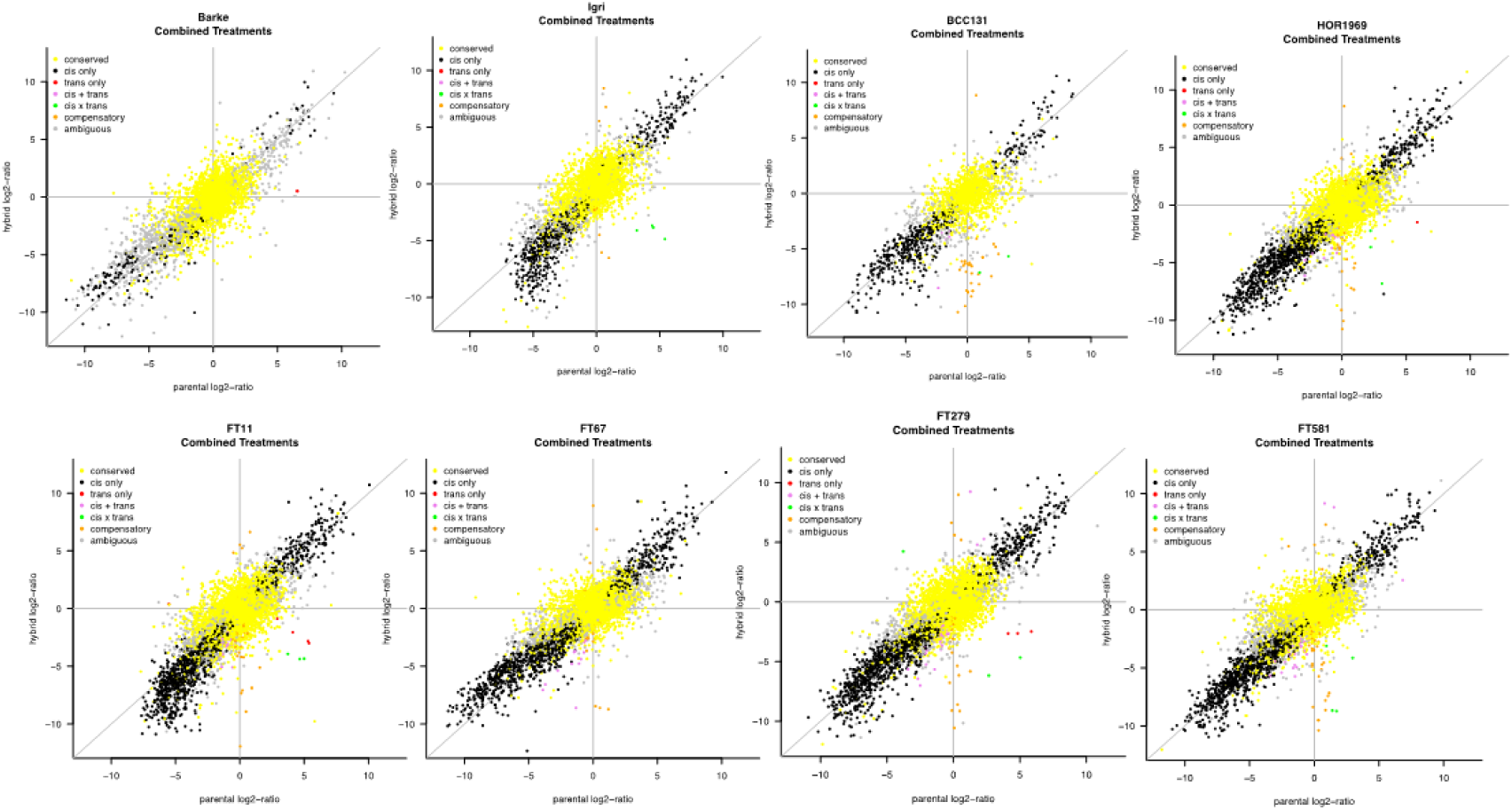
Log_2_ ratio plots of parents (x-axis) vs. parental alleles in the hybrid (y-axis) for all crosses when treatments were not considered separately and instead grouped as additional replicates.

---

**Table S1.** Accession numbers for individual accessions deposited into the European Nucleotide Archive (ENA). The table is sorted numerically according to <u>the ENA a</u>ccession number.

| Type | Accession | Unique Name |
| --- | --- | --- |
| Study | PRJEB29972 | ena-STUDY-IPK-Gatersleben-27-11-2018-08:58:04:359-462 |
| Sample | ERS2915967<br>(SAMEA5131586) | Sample_B_034 |
| Sample | ERS2915968<br>(SAMEA5131587) | Sample_B_105 |
| Sample | ERS2915969<br>(SAMEA5131588) | Sample_B_012 |
| Sample | ERS2915970<br>(SAMEA5131589) | Sample_B_125 |
| Sample | ERS2915971<br>(SAMEA5131590) | Sample_B_026 |
| Sample | ERS2915972<br>(SAMEA5131591) | Sample_B_117 |
| Sample | ERS2915973<br>(SAMEA5131592) | Sample_B_075 |
| Sample | ERS2915974<br>(SAMEA5131593) | Sample_B_065 |
| Sample | ERS2915975<br>(SAMEA5131594) | Sample_B_089 |
| Sample | ERS2915976<br>(SAMEA5131595) | Sample_B_001 |
| Sample | ERS2915977<br>(SAMEA5131596) | Sample_B_061 |
| Sample | ERS2915978<br>(SAMEA5131597) | Sample_B_027 |
| Sample | ERS2915979<br>(SAMEA5131598) | Sample_B_045 |
| Sample | ERS2915980<br>(SAMEA5131599) | Sample_B_112 |
| Sample | ERS2915981<br>(SAMEA5131600) | Sample_B_126 |
| Sample | ERS2915982<br>(SAMEA5131601) | Sample_B_092 |
| Sample | ERS2915983<br>(SAMEA5131602) | Sample_B_128 |
| Sample | ERS2915984<br>(SAMEA5131603) | Sample_B_056 |
| Sample | ERS2915985<br>(SAMEA5131604) | Sample_B_096 |
| Sample | ERS2915986<br>(SAMEA5131605) | Sample_B_031 |
| Sample | ERS2915987<br>(SAMEA5131606) | Sample_B_088 |
| Sample | ERS2915988<br>(SAMEA5131607) | Sample_B_110 |
| Sample | ERS2915989<br>(SAMEA5131608) | Sample_B_019 |
| Sample | ERS2915990<br>(SAMEA5131609) | Sample_B_082 |
| Sample | ERS2915991<br>(SAMEA5131610) | Sample_B_054 |
| Sample | ERS2915992<br>(SAMEA5131611) | Sample_B_050 |
| Sample | ERS2915993<br>(SAMEA5131612) | Sample_B_098 |
| Sample | ERS2915994<br>(SAMEA5131613) | Sample_B_039 |
| Sample | ERS2915995<br>(SAMEA5131614) | Sample_B_046 |
| Sample | ERS2915996<br>(SAMEA5131615) | Sample_B_017 |
| Sample | ERS2915997<br>(SAMEA5131616) | Sample_B_124 |
| Sample | ERS2915998<br>(SAMEA5131617) | Sample_B_129 |
| Sample | ERS2915999<br>(SAMEA5131618) | Sample_B_120 |
| Sample | ERS2916000<br>(SAMEA5131619) | Sample_B_006 |
| Sample | ERS2916001<br>(SAMEA5131620) | Sample_B_076 |
| Sample | ERS2916002<br>(SAMEA5131621) | Sample_B_090 |
| Sample | ERS2916003<br>(SAMEA5131622) | Sample_B_106 |
| Sample | ERS2916004<br>(SAMEA5131623) | Sample_B_004 |
| Sample | ERS2916005<br>(SAMEA5131624) | Sample_B_072 |
| Sample | ERS2916006<br>(SAMEA5131625) | Sample_B_025 |
| Sample | ERS2916007<br>(SAMEA5131626) | Sample_B_048 |
| Sample | ERS2916008<br>(SAMEA5131627) | Sample_B_018 |
| Sample | ERS2916009<br>(SAMEA5131628) | Sample_B_064 |
| Sample | ERS2916010<br>(SAMEA5131629) | Sample_B_033 |
| Sample | ERS2916011<br>(SAMEA5131630) | Sample_B_116 |
| Sample | ERS2916012<br>(SAMEA5131631) | Sample_B_058 |
| Sample | ERS2916013<br>(SAMEA5131632) | Sample_B_008 |
| Sample | ERS2916014<br>(SAMEA5131633) | Sample_B_084 |
| Sample | ERS2916015<br>(SAMEA5131634) | Sample_B_029 |
| Sample | ERS2916016<br>(SAMEA5131635) | Sample_B_086 |
| Sample | ERS2916017<br>(SAMEA5131636) | Sample_B_070 |
| Sample | ERS2916018<br>(SAMEA5131637) | Sample_B_114 |
| Sample | ERS2916019<br>(SAMEA5131638) | Sample_B_052 |
| Sample | ERS2916020<br>(SAMEA5131639) | Sample_B_100 |
| Sample | ERS2916021<br>(SAMEA5131640) | Sample_B_068 |
| Sample | ERS2916022<br>(SAMEA5131641) | Sample_B_122 |
| Sample | ERS2916023<br>(SAMEA5131642) | Sample_B_041 |
| Sample | ERS2916024<br>(SAMEA5131643) | Sample_B_080 |
| Sample | ERS2916025<br>(SAMEA5131644) | Sample_B_037 |
| Sample | ERS2916026<br>(SAMEA5131645) | Sample_B_104 |
| Sample | ERS2916027<br>(SAMEA5131646) | Sample_B_074 |
| Sample | ERS2916028<br>(SAMEA5131647) | Sample_B_002 |
| Sample | ERS2916029<br>(SAMEA5131648) | Sample_B_062 |
| Sample | ERS2916030<br>(SAMEA5131649) | Sample_B_044 |
| Sample | ERS2916031<br>(SAMEA5131650) | Sample_B_118 |
| Sample | ERS2916032<br>(SAMEA5131651) | Sample_B_013 |
| Sample | ERS2916033<br>(SAMEA5131652) | Sample_B_108 |
| Sample | ERS2916034<br>(SAMEA5131653) | Sample_B_016 |
| Sample | ERS2916035<br>(SAMEA5131654) | Sample_B_060 |
| Sample | ERS2916036<br>(SAMEA5131655) | Sample_B_094 |
| Sample | ERS2916037<br>(SAMEA5131656) | Sample_B_015 |
| Sample | ERS2916038<br>(SAMEA5131657) | Sample_B_035 |
| Sample | ERS2916039<br>(SAMEA5131658) | Sample_B_021 |
| Sample | ERS2916040<br>(SAMEA5131659) | Sample_B_066 |
| Sample | ERS2916041<br>(SAMEA5131660) | Sample_B_102 |
| Sample | ERS2916042<br>(SAMEA5131661) | Sample_B_078 |
| Sample | ERS2916043<br>(SAMEA5131667) | Sample_B_121 |
| Sample | ERS2916044<br>(SAMEA5131668) | Sample_B_053 |
| Sample | ERS2916045<br>(SAMEA5131669) | Sample_B_109 |
| Sample | ERS2916046<br>(SAMEA5131670) | Sample_B_107 |
| Sample | ERS2916047<br>(SAMEA5131671) | Sample_B_095 |
| Sample | ERS2916048<br>(SAMEA5131672) | Sample_B_007 |
| Sample | ERS2916049<br>(SAMEA5131673) | Sample_B_063 |
| Sample | ERS2916050<br>(SAMEA5131674) | Sample_B_057 |
| Sample | ERS2916051<br>(SAMEA5131675) | Sample_B_115 |
| Sample | ERS2916052<br>(SAMEA5131676) | Sample_B_099 |
| Sample | ERS2916053<br>(SAMEA5131677) | Sample_B_077 |
| Sample | ERS2916054<br>(SAMEA5131678) | Sample_B_079 |
| Sample | ERS2916055<br>(SAMEA5131679) | Sample_B_022 |
| Sample | ERS2916056<br>(SAMEA5131680) | Sample_B_123 |
| Sample | ERS2916057<br>(SAMEA5131681) | Sample_B_024 |
| Sample | ERS2916058<br>(SAMEA5131682) | Sample_B_055 |
| Sample | ERS2916059<br>(SAMEA5131683) | Sample_B_083 |
| Sample | ERS2916060<br>(SAMEA5131684) | Sample_B_038 |
| Sample | ERS2916061<br>(SAMEA5131685) | Sample_B_051 |
| Sample | ERS2916062<br>(SAMEA5131686) | Sample_B_003 |
| Sample | ERS2916063<br>(SAMEA5131687) | Sample_B_101 |
| Sample | ERS2916064<br>(SAMEA5131688) | Sample_B_093 |
| Sample | ERS2916065<br>(SAMEA5131689) | Sample_B_097 |
| Sample | ERS2916066<br>(SAMEA5131690) | Sample_B_127 |
| Sample | ERS2916067<br>(SAMEA5131691) | Sample_B_067 |
| Sample | ERS2916068<br>(SAMEA5131692) | Sample_B_059 |
| Sample | ERS2916069<br>(SAMEA5131693) | Sample_B_081 |
| Sample | ERS2916070<br>(SAMEA5131694) | Sample_B_043 |
| Sample | ERS2916071<br>(SAMEA5131695) | Sample_B_103 |
| Sample | ERS2916072<br>(SAMEA5131696) | Sample_B_005 |
| Sample | ERS2916073<br>(SAMEA5131697) | Sample_B_032 |
| Sample | ERS2916074<br>(SAMEA5131698) | Sample_B_111 |
| Sample | ERS2916075<br>(SAMEA5131699) | Sample_B_069 |
| Sample | ERS2916076<br>(SAMEA5131700) | Sample_B_085 |
| Sample | ERS2916077<br>(SAMEA5131701) | Sample_B_020 |
| Sample | ERS2916078<br>(SAMEA5131702) | Sample_B_073 |
| Sample | ERS2916079<br>(SAMEA5131703) | Sample_B_036 |
| Sample | ERS2916080<br>(SAMEA5131704) | Sample_B_119 |
| Sample | ERS2916081<br>(SAMEA5131705) | Sample_B_040 |
| Sample | ERS2916082<br>(SAMEA5131706) | Sample_B_023 |
| Sample | ERS2916083<br>(SAMEA5131707) | Sample_B_028 |
| Sample | ERS2916084<br>(SAMEA5131708) | Sample_B_113 |
| Sample | ERS2916085<br>(SAMEA5131709) | Sample_B_071 |
| Sample | ERS2916086<br>(SAMEA5131710) | Sample_B_087 |
| Sample | ERS2916087<br>(SAMEA5131711) | Sample_B_049 |
| Sample | ERS2916088<br>(SAMEA5131712) | Sample_B_047 |
| Sample | ERS2916089<br>(SAMEA5131713) | Sample_B_091 |
| Sample | ERS2916090<br>(SAMEA5131714) | Sample_B_030 |
| Sample | ERS2916091<br>(SAMEA5131715) | Sample_B_130 |
| Sample | ERS2916092<br>(SAMEA5131716) | Sample_B_042 |
| Sample | ERS2916093<br>(SAMEA5131717) | Sample_B_009 |

**Table S2.** Gene category assignment for Barke × Morex.

| Cold<br>Control | cis only | trans only | cis + trans | cis × trans | conserved | compensatory | ambiguous |
| --- | --- | --- | --- | --- | --- | --- | --- |
| cis only | 8 | 0 | 0 | 0 | 2 | 0 | 273 |
| trans only | 0 | 0 | 0 | 0 | 0 | 0 | 0 |
| cis + trans | 0 | 0 | 0 | 0 | 0 | 0 | 0 |
| cis × trans | 0 | 0 | 0 | 0 | 0 | 0 | 0 |
| conserved | 0 | 0 | 0 | 0 | 3,738 | 0 | 231 |
| compensatory | 0 | 0 | 0 | 0 | 0 | 0 | 0 |
| ambiguous | 0 | 0 | 0 | 0 | 220 | 0 | 454 |

**Table S3.** Gene category assignment for Igri × Morex.

| Cold<br>Control | cis only | trans only | cis + trans | cis × trans | conserved | compensatory | ambiguous |
| --- | --- | --- | --- | --- | --- | --- | --- |
| cis only | 172 | 0 | 0 | 0 | 34 | 0 | 134 |
| trans only | 0 | 0 | 0 | 0 | 0 | 0 | 0 |
| cis + trans | 0 | 0 | 0 | 0 | 0 | 0 | 0 |
| cis × trans | 0 | 0 | 0 | 1 | 0 | 0 | 0 |
| conserved | 18 | 0 | 0 | 0 | 3,713 | 0 | 193 |
| compensatory | 1 | 0 | 0 | 0 | 1 | 1 | 0 |
| ambiguous | 39 | 0 | 0 | 0 | 168 | 0 | 159 |

**Table S4.**
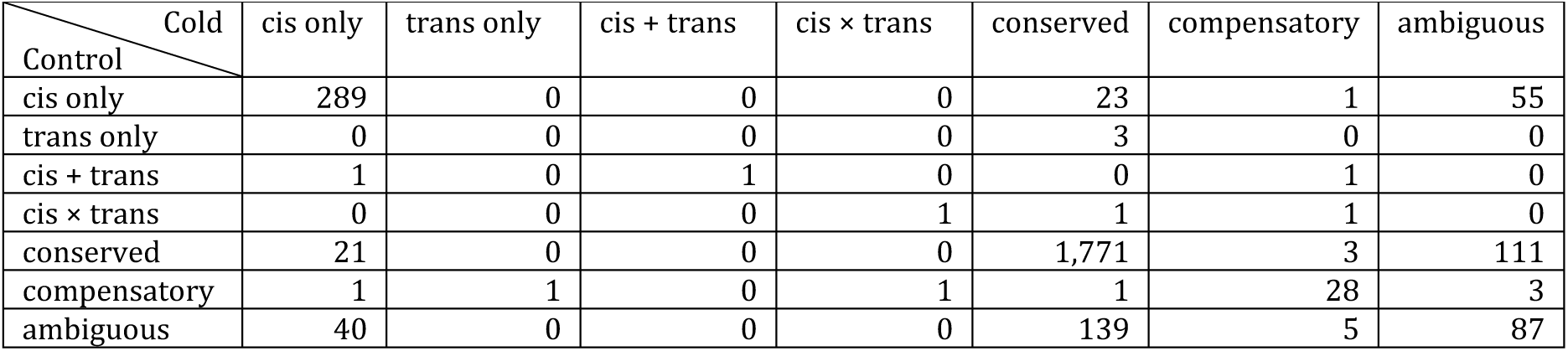
Gene category assignment for BCC131 × Morex.

| Cold<br>Control | cis only | trans only | cis + trans | cis × trans | conserved | compensatory | ambiguous |
| --- | --- | --- | --- | --- | --- | --- | --- |
| cis only | 289 | 0 | 0 | 0 | 23 | 1 | 55 |
| trans only | 0 | 0 | 0 | 0 | 3 | 0 | 0 |
| cis + trans | 1 | 0 | 1 | 0 | 0 | 1 | 0 |
| cis × trans | 0 | 0 | 0 | 1 | 1 | 1 | 0 |
| conserved | 21 | 0 | 0 | 0 | 1,771 | 3 | 111 |
| compensatory | 1 | 1 | 0 | 1 | 1 | 28 | 3 |
| ambiguous | 40 | 0 | 0 | 0 | 139 | 5 | 87 |

**Table S5.**
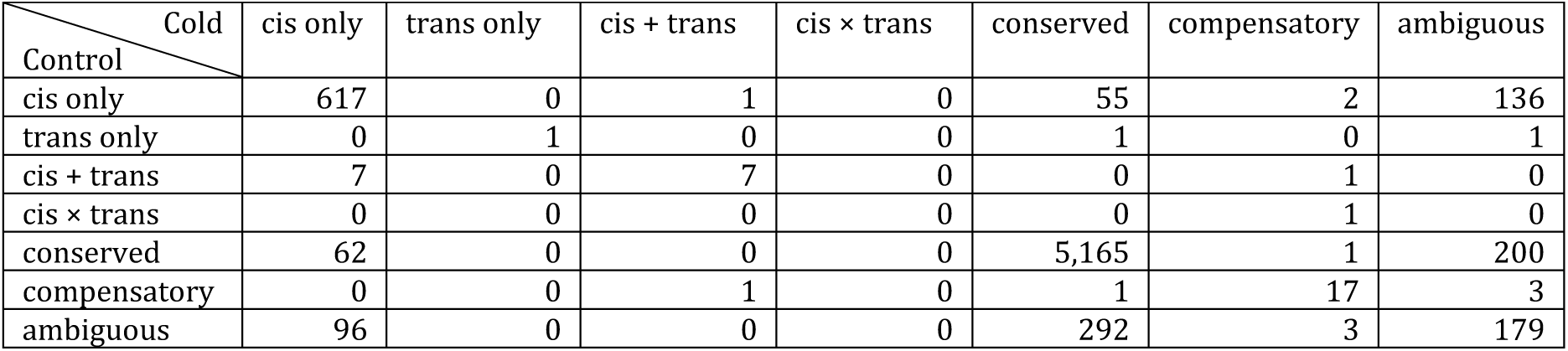
Gene category assignment for HOR1969 × Morex.

| Cold<br>Control | cis only | trans only | cis + trans | cis × trans | conserved | compensatory | ambiguous |
| --- | --- | --- | --- | --- | --- | --- | --- |
| cis only | 617 | 0 | 1 | 0 | 55 | 2 | 136 |
| trans only | 0 | 1 | 0 | 0 | 1 | 0 | 1 |
| cis + trans | 7 | 0 | 7 | 0 | 0 | 1 | 0 |
| cis × trans | 0 | 0 | 0 | 0 | 0 | 1 | 0 |
| conserved | 62 | 0 | 0 | 0 | 5,165 | 1 | 200 |
| compensatory | 0 | 0 | 1 | 0 | 1 | 17 | 3 |
| ambiguous | 96 | 0 | 0 | 0 | 292 | 3 | 179 |

**Table S6.**
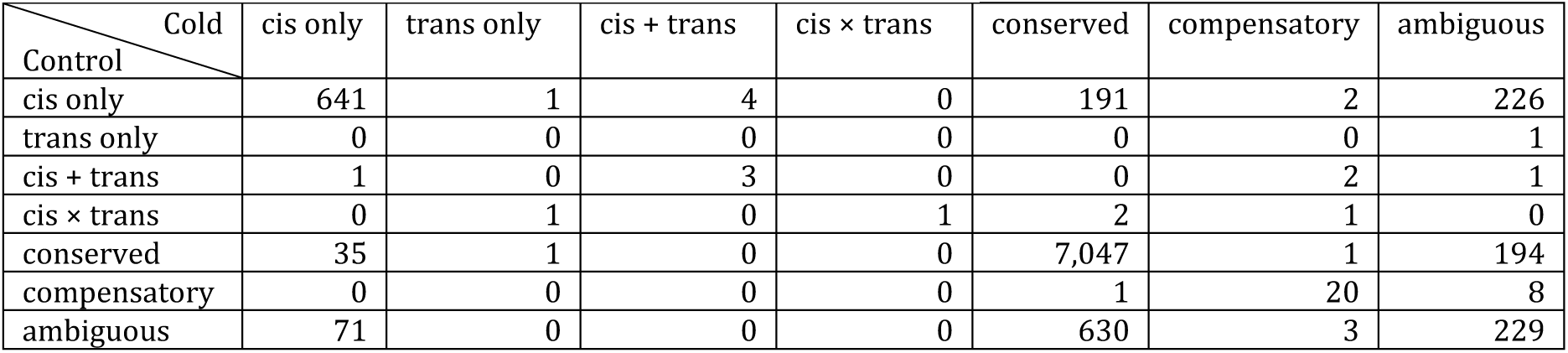
Gene category assignment for FT11 × Morex.

**Table S7.** Gene category assignment for FT67 × Morex.

| Cold<br>Control | cis only | trans only | cis + trans | cis × trans | conserved | compensatory | ambiguous |
| --- | --- | --- | --- | --- | --- | --- | --- |
| cis only | 754 | 0 | 5 | 0 | 51 | 1 | 151 |
| trans only | 1 | 0 | 0 | 0 | 2 | 0 | 0 |
| cis + trans | 3 | 0 | 9 | 0 | 0 | 2 | 0 |
| cis × trans | 0 | 0 | 0 | 0 | 0 | 0 | 0 |
| conserved | 41 | 0 | 0 | 0 | 6,408 | 0 | 255 |
| compensatory | 0 | 0 | 0 | 0 | 0 | 12 | 5 |
| ambiguous | 105 | 0 | 0 | 0 | 484 | 1 | 300 |

**Table S8.** Gene category assignment for FT279 × Morex.

| Cold<br>Control | cis only | trans only | cis + trans | cis × trans | conserved | compensatory | ambiguous |
| --- | --- | --- | --- | --- | --- | --- | --- |
| cis only | 749 | 0 | 5 | 0 | 45 | 2 | 93 |
| trans only | 0 | 0 | 0 | 1 | 1 | 0 | 0 |
| cis + trans | 3 | 0 | 12 | 0 | 0 | 0 | 0 |
| cis × trans | 0 | 0 | 0 | 0 | 1 | 2 | 0 |
| conserved | 78 | 0 | 0 | 0 | 6,241 | 2 | 289 |
| compensatory | 0 | 0 | 1 | 0 | 2 | 19 | 6 |
| ambiguous | 137 | 1 | 0 | 1 | 365 | 4 | 222 |

**Table S9.** Gene category assignment for FT581 × Morex.

| Cold<br>Control | cis only | trans only | cis + trans | cis × trans | conserved | compensatory | ambiguous |
| --- | --- | --- | --- | --- | --- | --- | --- |
| cis only | 789 | 0 | 9 | 0 | 43 | 0 | 111 |
| trans only | 2 | 0 | 0 | 0 | 1 | 0 | 1 |
| cis + trans | 4 | 0 | 6 | 0 | 0 | 2 | 0 |
| cis × trans | 0 | 0 | 0 | 2 | 1 | 4 | 2 |
| conserved | 81 | 1 | 0 | 0 | 6,582 | 0 | 360 |
| compensatory | 0 | 0 | 3 | 1 | 1 | 20 | 4 |
| ambiguous | 157 | 0 | 0 | 1 | 465 | 3 | 283 |

